# Extracellular GPX3 drives insulin receptor expression in adipose tissue independent of its peroxidase activity

**DOI:** 10.64898/2026.10.02.755889

**Authors:** Robert Hauffe, Marina González, Simran Chopra, J. Sophie Messner, Christoph Schüßler, Maria Maares, Vanessa Kabbouchy, Attila Andor, Elias Arnér, André Kleinridders

**Author notes:** Address correspondence to: André Kleinridders, Department of Molecular and Experimental Nutritional Medicine Institute of Nutritional Science, University of Potsdam, Arthur-Scheunert-Allee 114-116, 14558 Nuthetal, Germany.

## Abstract

Obesity-associated insulin resistance is driven in part by adipose tissue dysfunction and oxidative stress, yet the role of extracellular mechanisms in modulating insulin signaling remains poorly understood. Here, we show that extracellular Glutathione Peroxidase 3 (GPX3) enhances adipocyte insulin sensitivity and functionalities independent of its canonical peroxidase function. GPX3, a secreted selenoprotein highly expressed in white adipose tissue (WAT), is suppressed in obesity, but whether it actively regulates insulin sensitivity had not been established. We find that WAT *GPX3* expression closely mirrors insulin receptor (InsR) expression across a broad range of metabolic parameters in humans and increases dynamically following weight loss. Prolonged therapeutic selenium supplementation in diet-induced obese mice selectively suppressed epididymal WAT oxidative damage, restored GPX3 and InsR expression, and enhanced insulin-stimulated AKT phosphorylation, improving oral glucose tolerance. Mechanistically, recombinant GPX3 extracellularly administered at physiological concentrations induced InsR expression across multiple adipocyte models, including pre-adipocytes, epididymal WAT explants, and adipocyte organoids, by promoting binding of the transcription factor Specificity Protein 1 (SP1) to the *Insr* promoter, while knockdown of GPX3 completely blocks SVF adipogenesis. These effects rely on the structural integrity, rather than selenocysteine-dependent activity. Thus, we identify a GPX3-SP1-InsR signaling axis in adipose tissue insulin sensitivity, suggesting GPX3 is a novel paracrine mediator of metabolic health and a potential therapeutic target to reinstate WAT insulin sensitivity in obesity and type 2 diabetes.

**Graphical Abstract:** 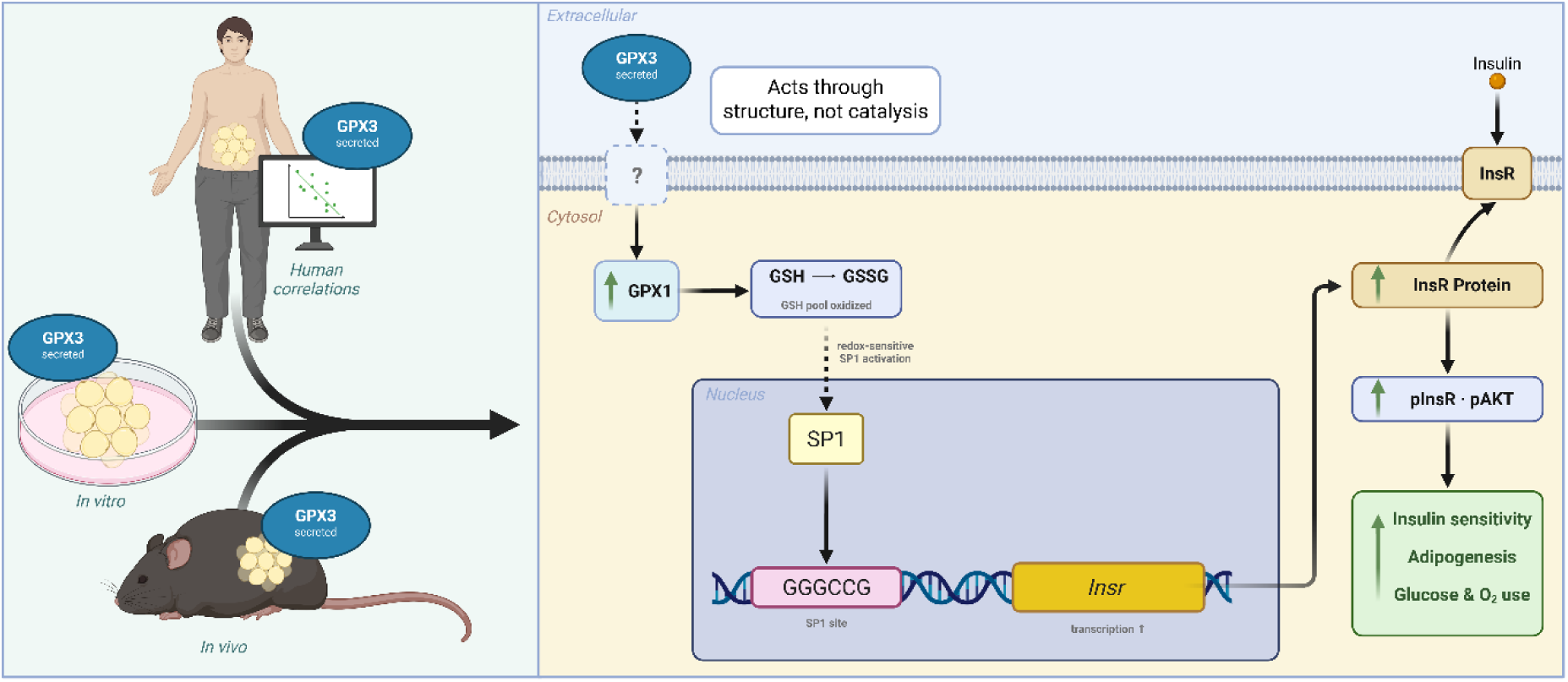

## Introduction

Insulin resistance is a defining feature of obesity-associated metabolic disease and a key driver of type 2 diabetes (T2D), cardiovascular disease, and metabolic dysfunction-associated steatotic liver disease (MASLD). While insulin resistance manifests systemically, white adipose tissue (WAT) plays a central initiating and amplifying role through its capacity to buffer lipid flux, regulate inflammation, and secrete endocrine factors that modulate whole-body insulin sensitivity (Kahn et al., 2006; Lyu et al., 2021; Scherer, 2019). Failure of adipose tissue function by oxidative stress and chronic low-grade inflammation leads to adipocyte hypertrophy, which impairs insulin receptor (INSR) signaling locally and in peripheral tissues (Guilherme et al., 2008).

Beyond its canonical role in lipid storage, insulation, and mechanical organ protection, adipose tissue is known as a major site of metabolic regulation and activity. A natural, and indeed necessary by-product of metabolic activity is the production of reactive oxygen species (ROS). Moderate levels of ROS and their reactive products are important for cellular metabolism. They fulfil signaling roles to inform the cells about the state of mitochondrial load and regulate adipocyte differentiation, insulin signaling, and transcriptional programs involved in metabolic homeostasis (Hoehn et al., 2009; Rhee, 2006). Insulin signaling itself is intrinsically redox-sensitive: transient increases in hydrogen peroxide (H₂O₂) enhance signaling by reversibly inhibiting protein tyrosine phosphatases such as PTP1B and PTEN, thereby prolonging INSR and AKT phosphorylation (Goldstein et al., 2005; Loh et al., 2009; May and de Haën, 1979). However, excessive or spatially dysregulated ROS accumulation promotes oxidative damage, impairs insulin action, and contributes to adipose tissue dysfunction (Furukawa et al., 2017).

When present in excess, e.g., by a temporary increase in respiration due to an increase in nutrient influx, ROS are detoxified through intracellular anti-oxidative systems. Remarkably, both the excessive presence, as well as forced absence (through exogenous antioxidant treatment) is decreasing adipocyte insulin sensitivity and subsequent functionality (Fernando et al., 2020).

To maintain this delicate balance, insulin-responsive tissues rely on tightly controlled antioxidant systems. While intracellular antioxidant enzymes, including catalase, peroxiredoxins, and glutathione peroxidases (GPX) have been studied extensively in the context of insulin resistance (Merry et al., 2014; Brigelius-Flohé and Flohé, 2020), comparatively little is known about how extracellular redox-related mechanisms may influence INSR signaling. This represents a critical gap in our understanding, as the INSR is a transmembrane protein whose activation and expression may be modulated by redox events occurring at or near the cell surface by unknown mechanisms.

Glutathione peroxidase 3 (GPX3) is a secreted selenoprotein that catalyzes the reduction of H₂O₂ and lipid hydroperoxides using glutathione as an electron donor. Unlike GPX1, which functions intracellularly, GPX3 is synthesized with an N-terminal signal peptide, secreted into the extracellular space, and circulates in plasma at relatively high concentrations, contributing around 10 – 15 % of circulating selenium (Brigelius-Flohé, 1999; Burk and Hill, 2015; Takahashi et al., 1987). GPX3 is highly expressed in kidney, adipose tissue, and lung tissue, and its circulating levels are sensitive to selenium availability, inflammatory cues, and metabolic status (Kerr et al., 2020; Langhardt et al., 2018; Lee et al., 2008; Zhang et al., 2024). Recently, proximity-dependent biotinylation studies have shown that GPX3 is actively secreted by white adipocytes, placing it firmly as an adipokine. Moreover, WAT-derived-GPX3 reaches the circulation and decreases significantly in high-fat conditions in mice, suggesting GPX3 levels react dynamically to changes in adiposity (Plucińska et al., 2026).

Several clinical studies have reported altered GPX3 expression or activity in obesity and type 2 diabetes, yet the directionality and interpretation of these associations have remained inconsistent (Baez-Duarte et al., 2014; Goyal et al., 2011; Langhardt et al., 2018). Importantly, most mechanistic work on selenium and GPX proteins in metabolism has focused on intracellular enzymes, particularly GPX1, whose overexpression can paradoxically promote insulin resistance by blunting physiological ROS signaling (Loh et al., 2009; McClung et al., 2004). In contrast to intracellular GPX1, the downregulation of GPX3 in the adipose tissue of obese individuals is of particularly interest given its high basal expression in healthy WAT (Lee et al., 2008). Clinical studies have shown that adipose GPX3 expression is suppressed by pro-inflammatory signals like TNFα and by the hypoxic conditions that arise when adipocyte size exceeds the oxygen diffusion limit (Fleischmann et al., 2005; Lee et al., 2008). This suppression creates a vicious cycle where reduced antioxidant capacity further exacerbates local oxidative stress, leading to adipocytokine dysregulation and systemic metabolic complications. Whether extracellular GPX3 exerts distinct effects on insulin sensitivity through mechanisms separate from intracellular antioxidant pathways is not known. We have shown previously that GPX3 is associated with metabolic health in both mice and humans, that its depletion decreases insulin sensitivity in vitro and is associated with the reduction of WAT INSR expression seen with increasing BMI (Hauffe et al., 2020, 2022).

Here, we test the hypothesis that GPX3 represents a redox-regulatory node linking adipose tissue function to INSR expression and signaling. By integrating human adipose tissue transcriptomic analyses, prolonged therapeutic selenite supplementation in obese mice, and a series of cell-based, ex vivo, and organoid models, we identify a GPX3-SP1-InsR axis that enhances adipocyte insulin responsiveness. Our findings uncover a previously unrecognized role for extracellular GPX in metabolic regulation and suggest that adipose-derived GPX3 contributes to systemic insulin sensitivity by fine-tuning signaling at the INSR.

## Results

### GPX3 expression associates with metabolic health in humans

To systematically assess the association of WAT-derived GPX3 and metabolic health we explored data from the adiposetissue.org repository (Zhong et al., 2025), which is a publicly available database that integrates multiple clinical and molecular datasets from human adipose in a manner akin to a meta-analysis. Here, *GPX3* expression in subcutaneous white adipose tissue (scWAT) correlates positively with markers of healthy adipose tissue functions like isoproterenol-stimulated lipolysis rates, and circulating HDL (Fig. 1A). Furthermore, WAT *GPX3* expression shows significant negative correlation with reported markers of deteriorating metabolic states. Namely, in order of descending correlation strength: fat cell volume, WAT leptin secretion, homeostatic model assessment of insulin resistance (HOMA-IR), circulating insulin, BMI, hip and waist circumference, waist-to-hip ratio, circulating triglycerides and C-reactive protein, HbA1c, and circulating glucose (Fig. 1A). This is specific to *GPX3*, as the main intracellular glutathione peroxidase, *GPX1*, shows the opposite correlations, where increases in the same aforementioned negative metabolic traits are associated with higher *GPX1* expression in the adipose (Suppl. Fig. S1A), with the only exception of circulating glucose, which is not correlated with WAT *GPX1* expression. Expression of *TXNRD1*, another gene encoding a major selenoprotein previously shown to suppress INSR responsiveness by intracellular mechanisms (Peng et al., 2016), showed a similar inverse picture, with positive correlations to beneficial metabolic traits, and with negative correlations to unhealthy metabolic traits (Suppl. Fig. S1B), suggesting a dysregulation of GPX3 in overweight and obese conditions. As mentioned, INSR expression negatively correlates with body weight or BMI (Hauffe et al., 2020, 2022). Moreover, upon comparing the correlation calculations between clinical parameters and the expression of *GPX3* or *INSR*, there is a strong overlap in both the direction and magnitude of these associations (Fig. 1B) indicating that GPX3 plays a crucial role for adipose tissue insulin sensitivity. Due to the connection between altered bodyweight and both WAT-specific and circulating GPX3 levels, we next explored the datasets for weight loss interventions within adiposetissue.org. Here, data from both surgical and dietary interventions (Suppl. Fig. S1C,D) reveal that WAT-derived *GPX3* expression increases after weight loss, and is not restricted to a specific weight loss intervention (Fig. 1C, Suppl. Fig. S1E). Interestingly, this seems to operate on a dynamic scale, with GPX3 expression increasing in proportion to the extent of weight loss. While moderate changes are observed during shorter dietary interventions (5 and 12 weeks), larger effects are seen with longer interventions (14 weeks) and the largest effects are seen following substantial weight loss after bariatric surgery. This is also apparent for *INSR* expression in WAT, where shorter dietary interventions (5 and 12 weeks) are insufficient to increase INSR expression, longer dietary intervention (14 weeks) leads to a significant *INSR* increase, and bariatric surgery results in the largest increase in *INSR* levels (Suppl. Fig. S1F). Together, these findings suggest a dynamic regulation where changes in GPX3 occur relatively rapidly after weight loss, while INSR expression increases more gradually.

**Figure 1:**
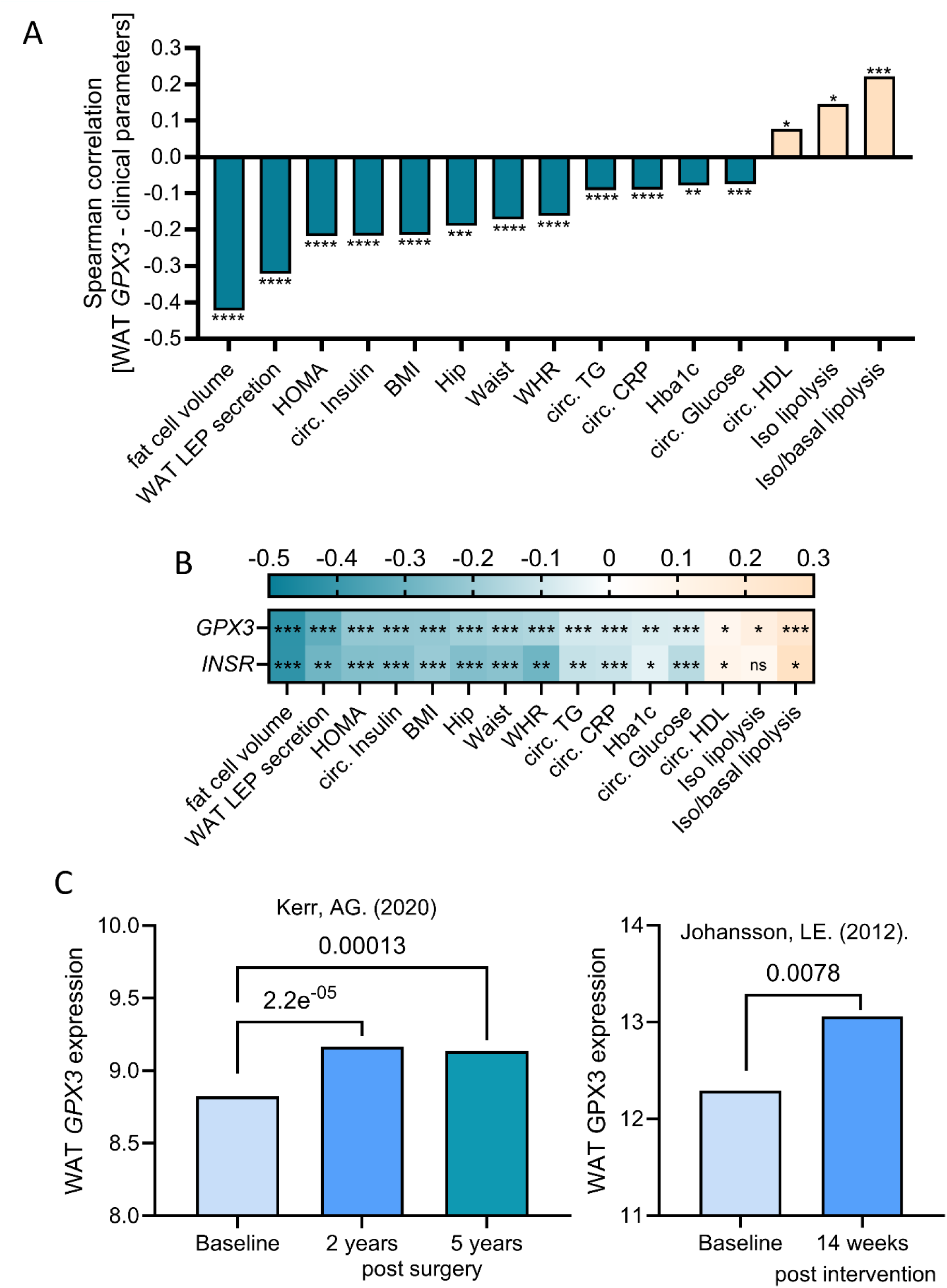
GPX3 expression associates with metabolic health in humans. **(A)** Correlation of WAT *GPX3* expression and metabolic traits. Bar height represents Spearman correlation coefficient (ρ). Asterisks indicate -log₁₀(p) category (* p ≥ 10-20, ** p ≥ 20–30, *** p ≥ 30–60, **** p > 60 **(B)** Heatmap comparing correlations of metabolic traits with WAT *GPX3* and *INSR* expression. Color scale indicates the level of correlation and FDR-adjusted significance is indicated by asterisks (*** p_FDR_ <0.001, ** p_FDR_ < 0.01, * p_FDR_ < 0.05). **(C)** WAT *GPX3* expression increases after weight loss, through either bariatric surgery or diet. Statistical differences between baseline and follow-up time points were assessed using the Wilcoxon signed-rank test, as implemented in adiposetissue.org. All data extracted from adiposetissue.org v1.0.0, see method section for references.

### Therapeutic selenite slows body weight gain

Considering this relatively slower response of the INSR in dietary interventions, we concluded that a likely factor for a lack of improved insulin sensitivity or body weight in our previously published therapeutic selenite study, was inadequate intervention time to impact adipose tissue health (Hauffe et al., 2022). Therefore, we extended the time mice were fed the selenite rich high-fat diet (SRHFD) to twice the original time frame (20 weeks vs. 10 weeks, see Fig. 2A). This intervention regime was sufficient to impact body weight, with a 20 % decrease in body weight gain at the end of the study compared to selenium adequate HFD-fed mice (Fig. 2B). This was mainly due to a decrease in fat, as seen by a significant reduction in epididymal and subcutaneous white adipose tissue weight (eWAT, scWAT, Fig. 2C) for a total reduction of 13 % WAT weight when combining both depots (Fig. 2D). We also found a significant reduction in kidney weight (-12 %, Fig. 2C). The reduction in WAT mass was also associated with a 70 % increase in average epididymal adipocyte number, a 44 % decreased average adipocyte area, and consequently, smaller adipocyte circumference (28 % decrease, Fig. 2E-H), all indicating a decrease in adipocyte hypertrophy. Interestingly, analyzing selenium content in various tissues revealed increased levels specifically in the liver, kidney, and scWAT, but not in the eWAT (Fig. 2I), suggesting depot-specific regulation of selenium supply. To gain insights into the differences of the metabolic phenotype between our published preventive study, where mice were fed a selenite-rich diet before onset of obesity, the short-term intervention, and our new long-term intervention, we assessed total selenium content in tissues. This analysis revealed that selenium content in tissues was only slightly, but significantly, elevated by 8 % in the short-term supplementation (10 weeks), which increased to 24 % higher selenium content in the 20-week supplementation after established obesity (Fig. 2J). Of note, supplementing before establishing obesity (preventive study, Hauffe et al., 2020) increases selenium content by 38 %, suggesting a selenium uptake impairment in obese conditions. This effect does not seem to be an overall improvement in trace element uptake and assimilation, as copper, iron, and zinc were not found in higher quantities (Suppl. Fig. S2A). However, we did observe a 21 % increase in manganese levels across tissues. As the diets do not differ in manganese content (Hauffe et al., 2020), the reason for this remains to be elucidated.

**Figure 2:**
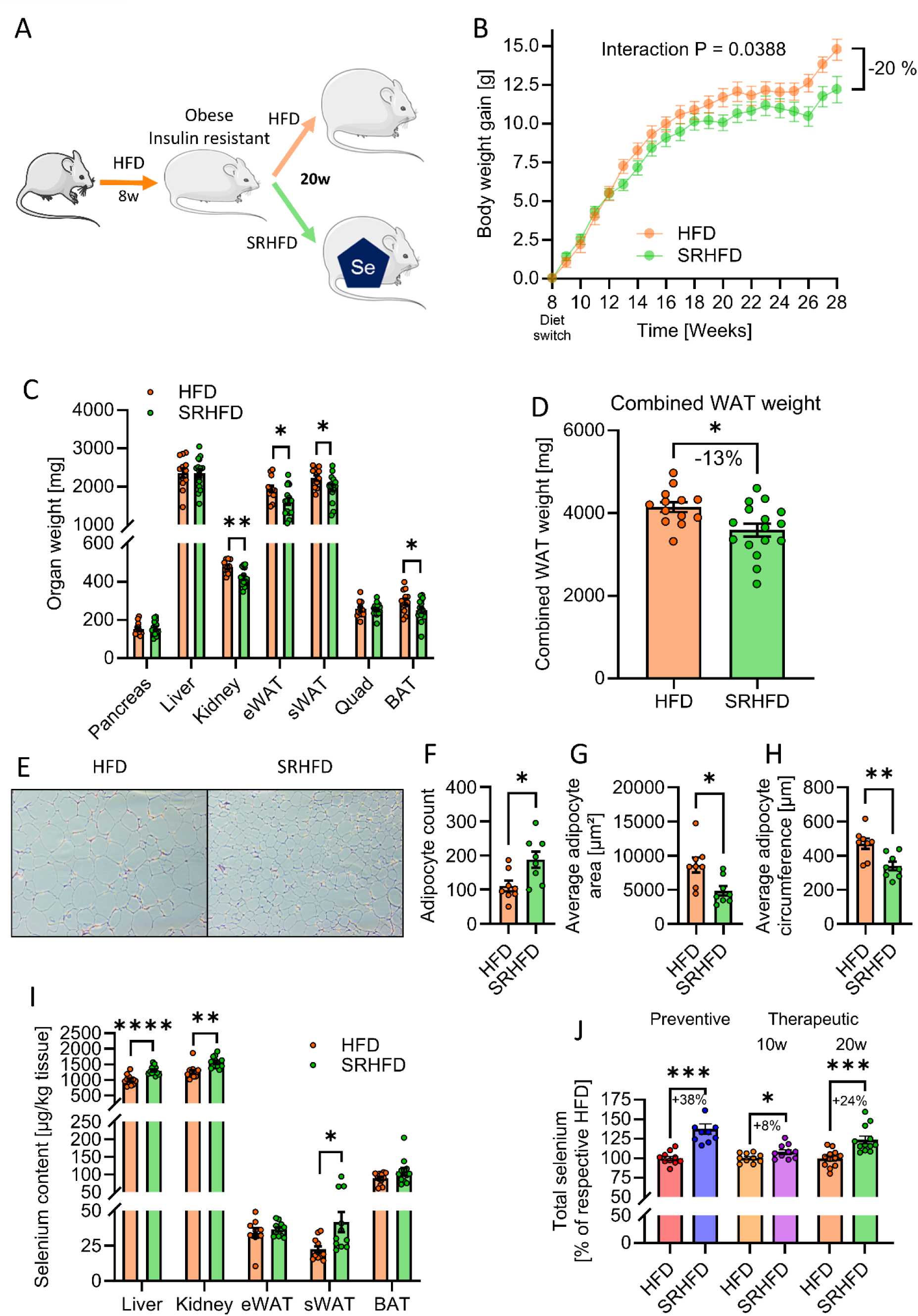
Mice fed a selenium-rich high fat diet after established obesity show decreased weight gain and fat accumulation. All data in B-J compares male C57BL/6N mice fed either HFD or SRHFD after established obesity for 20 additional weeks. **(A)** Experimental setup, HFD = High fat diet, 60 % calories from fat and ∼230 ng Se/g diet; SRHFD = Selenium rich HFD, ∼670 ng Se/g diet, w = weeks **(B)** Body weight gain (N=20/group). **(C)** Organ weights (N=14/group). **(D)** Combined white adipose tissue weights (recreated from C, (N=14/group). **(E-H)** representative microscopy image (E) and calculated number (F), area (G), and circumference (H) of eWAT cells (N=8/group). **(I)** Tissue selenium content (N=12/group). **(J)** Total tissue selenium contents of male C57BL/6N mice fed either HFD or SRHFD at the start of the intervention (preventive, recreated from Hauffe et al 2020) or for 10 weeks (recreated from Hauffe et al 2022) and 20 weeks, respectively, after established obesity, 20w data aggregated from panel I as average selenium per mouse). All data are presented as mean ± SEM. Data in B was analyzed by a Mixed-effects model with the Geisser-Greenhouse correction and Šidák’s multiple comparisons test. Significances in C-J after 2-tailed Welch’s t-test, *: p< 0.05, **: p< 0.01, ***: p< 0.001, ****: p< 0.0001.

### Adipose tissue redox state and insulin sensitivity is improved after selenite supplementation

Given the high expression of the selenoprotein GPX3 in adipose tissue, we next examined GPX expression levels in adipose depots and its association with InsR expression. In line with our hypothesis of beneficial effects in prolonged treatment, InsR expression in both scWAT and eWAT was increased by 29 % and 192 %, respectively, relative to HFD-fed mice (Suppl. Fig. S2B, Fig. 3A). Additionally, GPX1 was increased by 88 % and 82 %, whereas GPX3 showed a 59 % increase in scWAT, but a drastic 466 % increase in eWAT (Suppl. Fig. S2B, Fig. 3A). This highlights a disconnection between selenium supply and specific protein regulation patterns, as the scWAT, but not eWAT, showed an elevated selenium content (Fig. 2I). This disconnect continues across other tissues. In liver and kidney, where selenium content was significantly increased, only the kidney shows elevated GPX1 and GPX3 expression, which may reflect their individual roles in selenium metabolism. Interestingly, InsR expression was unaltered in both of these tissues (Suppl. Fig. S2B). Elevated GPX proteins in WAT should alter intracellular redox states and, thus, antioxidant capacity. Indeed, performing a GPX activity assay on eWAT samples, using H_2_O_2_ and reduced glutathione as substrates, showed a markedly steeper NADPH depletion in the selenite-treated group, indicating a more pronounced glutathione oxidation by intracellular glutathione peroxidases in eWAT from SRHFD-fed mice (70 % increase in area under the curve, Fig. 3B). Of note, this assay cannot differentiate between different GPX proteins. We speculated whether this increased antioxidative capacity could counteract obesity-induced oxidative damage. Consequently, when we assessed oxidative damage by blotting for protein carbonylation in eWAT, we observed a significant decrease of protein carbonyls in the SRHFD group compared to control (Fig. 3C). Furthermore, as oxidative stress can induce insulin resistance, we performed vena cava insulin stimulation to analyze tissue-specific insulin sensitivity. As suggested by elevated InsR expression, downstream activation of AKT at serine 473 was significantly enhanced by 76 % in the SRHFD group in eWAT (Fig. 3D). Crucially, this insulin sensitizing effect was specific to the eWAT, as neither the scWAT, brown adipose tissue (BAT), nor other insulin-sensitive tissues, such as the liver and skeletal muscle showed differences in AKT phosphorylation (Suppl. Fig. S3A). This did not lead to marked differences in systemic insulin sensitivity, as seen by identical glucose curves during an ITT (Suppl. Fig. S3B). Intriguingly, and consistent with improved eWAT insulin sensitivity, SRHFD mice demonstrated a blunted early glycemic excursion during oGTT, reflected by a reduced peak above baseline (-20 %) and a lower rate of glucose rise (ROC up, -22 %). The rate of subsequent glucose clearance was not significantly different between groups (ROC down), suggesting that the attenuation of peak glucose was driven primarily by differences in the early absorptive phase rather than enhanced systemic clearance capacity (Fig. 3E-G, Suppl. Fig. S3C). Furthermore, data from the International Mouse Phenotyping Consortium (IMPC, https://www.mousephenotype.org/, Wilson et al., 2026) for heterozygous GPX3 knockout mice (www.mousephenotype.org/data/genes/MGI:105102) shows impaired glucose tolerance (Suppl. Fig. S3D), suggesting a modulating role of GPX3 in whole-body glucose metabolism with adequate selenite supply. Taken together, prolonged selenite supplementation in established obesity increases GPX3 expression, counteracts oxidative damage and enhances epididymal adipose tissue insulin signaling, contributing to improved glucose tolerance.

**Figure 3:**
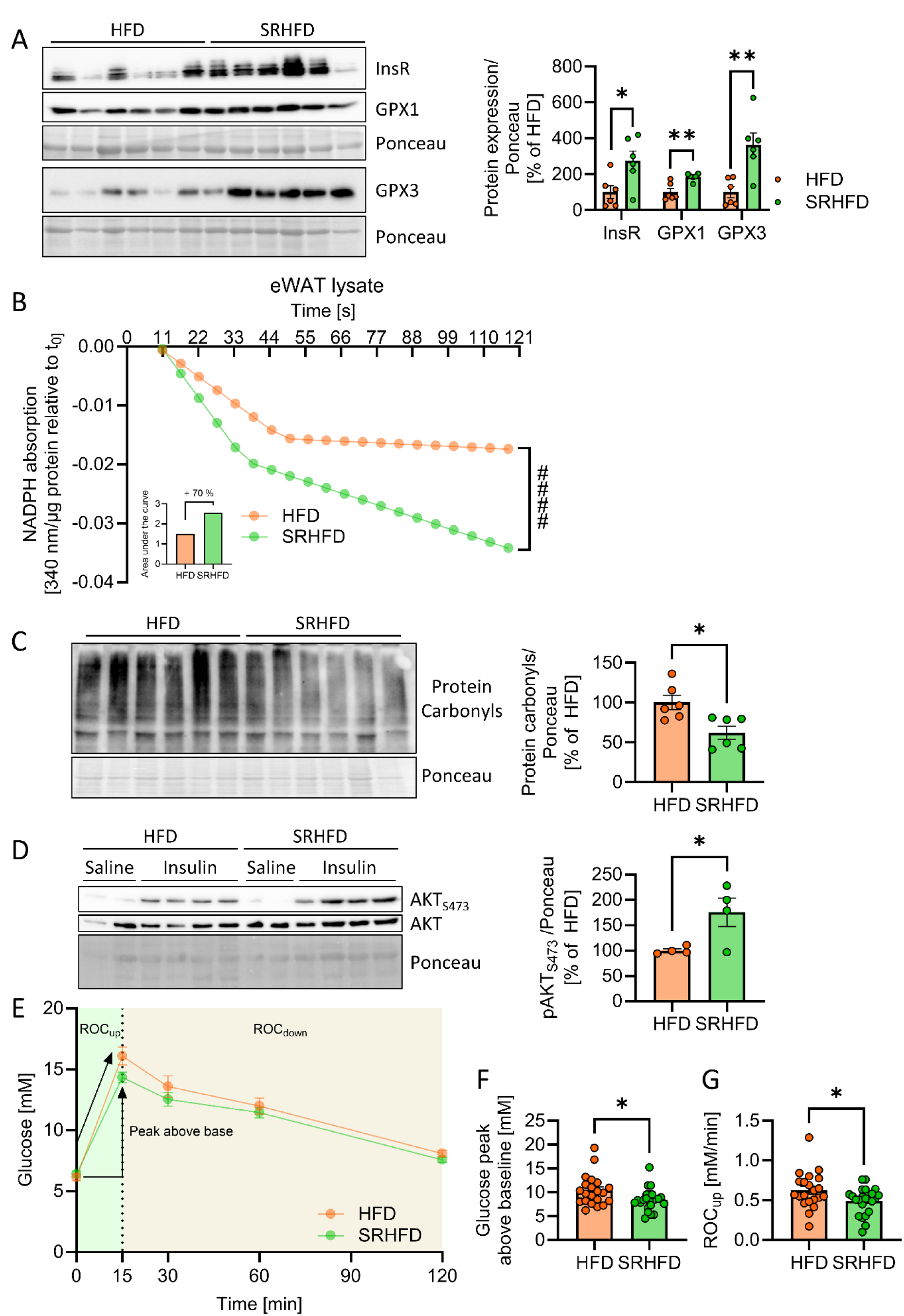
Epididymal WAT of mice fed a selenium-rich high fat diet after established obesity show decreased oxidative stress and improved insulin signaling. All data in A-D compares male C57BL/6N mice fed either HFD or SRHFD after established obesity for 20 additional weeks **(A)** Protein expression and densitometric analysis of insulin receptor and glutathione peroxidase 1 and 3 in eWAT. **(B)** GPX activity assay from eWAT lysates, decrease in NADPH absorption signifies activity (N=6/group). **(C)** Protein carbonylation in the eWAT detected via Western Blot and densitometric analysis. **(D)** Protein phosphorylation, expression and densitometric analysis of AKT in eWAT lysates after vena cava injection of saline or insulin. **(E-G)** Glucose curve (E), peak above baseline (F), and Rate of change from baseline to peak (ROC_up_, G) during an oGTT in mice fed either HFD or SRHFD for 16 weeks after established obesity (N=20/group). All data are presented as mean ± SEM. Data in B was analyzed by two-way ANOVA and Šidák’s multiple comparisons test. Significances in A, C, D, F and G after 2-tailed Welch’s t-test, *: p< 0.05, **: p< 0.01.

### Extracellular GPX3 induces Insr via SP1 independently of peroxidase activity

We have previously shown that selenite treatment increases extracellular GPX3 levels which correlate with increased InsR expression and signaling, indicating that the beneficial effects of our selenite supplementation may be due to elevated GPX3 levels (Hauffe et al., 2020, 2022). Within the GPX family GPX3 is unique in being the only secreted protein, evidenced by a highly conserved N-terminal secretion signal, that is not present in other members of the family (Suppl. Fig. S4A,B). Thus, the main GPX3 functions should likely be in the extracellular compartment.

To ensure delivery of catalytically active, extracellular GPX3, we generated murine recombinant GPX3 (rGPX3) using previously established protocols (Lajoie et al., 2013; Cheng and Arnér, 2017), showing dose-dependent peroxidase activity (Suppl. Fig. S5A). Of note, it has been shown that mature-adipocyte conditioned media shows GPX3 concentrations around 500 ng/ml (Lee et al., 2008). Therefore, treating cells with 500 ng/ml rGPX3 via addition to the media represents a physiologically relevant experimental setup *in vitro*. As a redox-reactive protein, we hypothesized that it is the peroxidase function of GPX3 that mediates any effects. To test this, we also generated a selenocysteine-to-cysteine mutant U49C of GPX3 (rGPX3^U49C^), which is devoid of GPX activity and enzymatically indistinguishable to heat-inactivated rGPx3 (HI-rGPX3, Suppl. Fig. S5B).

Surprisingly, treating 3T3-L1 pre-adipocytes with 500 ng/ml of either rGPX3 or rGPX3^U49C^ for 24 hours was sufficient to drive an increase in InsR expression to almost identical levels (Fig. 4A).

**Figure 4:**
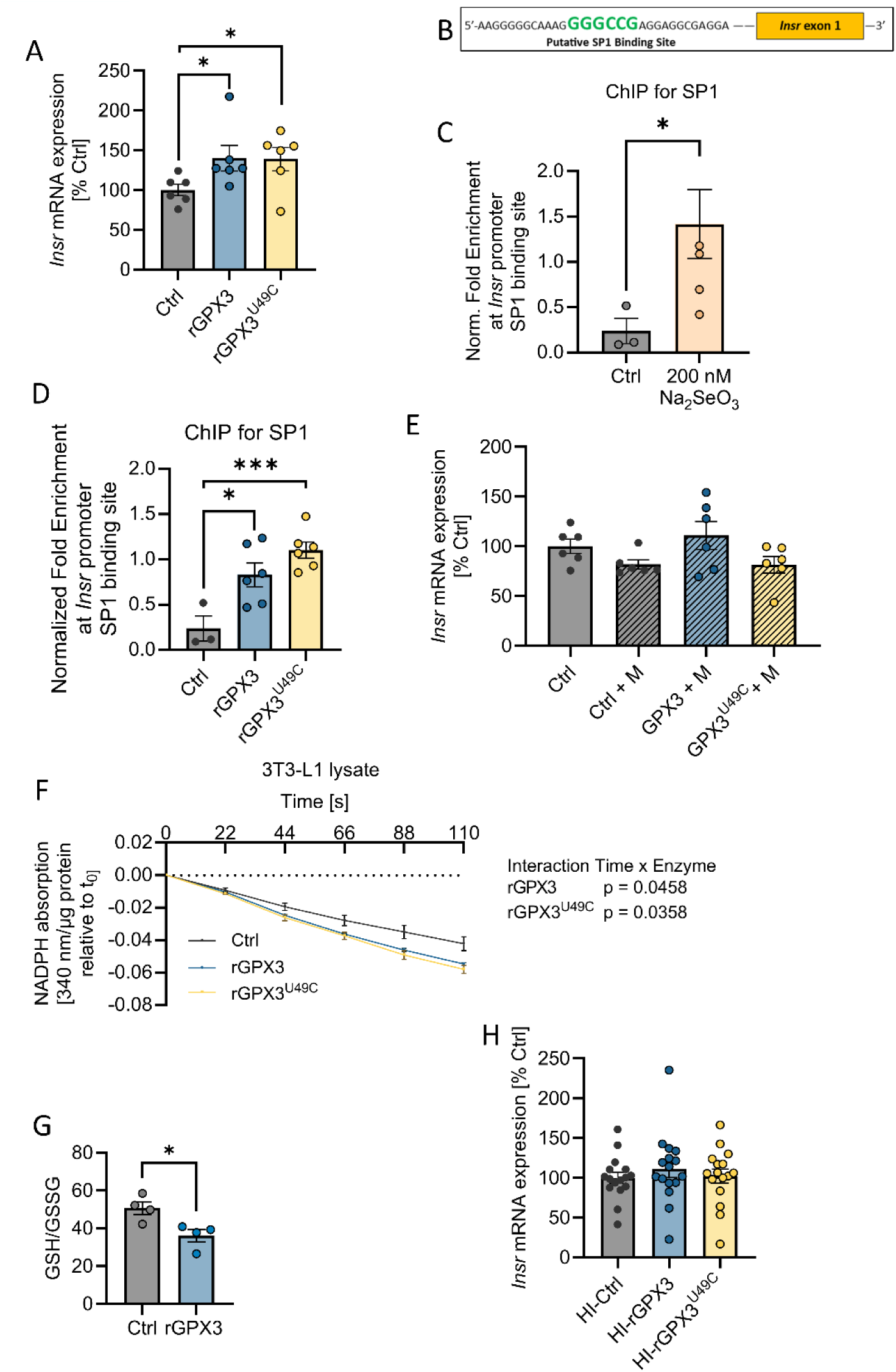
Extracellular recombinant GPX3 alters Insr expression via intracellular redox changes and SP1 activity independent of peroxidase function. **(A)** Gene expression of *Insr* in 3T3-L1 pre-adipocytes after treatment with functional (rGPX3) or selenocysteine-to-cysteine mutated enzyme (rGPX3^U49C^) for 24 h at 500 ng/ml. **(B)** Sequence within the mouse *Insr* promoter region showing putative SP1 binding site. **(C,D)** Normalized fold enrichment of SP1 to the putative *Insr* promoter binding site shown in B after sodium selenite (C, 24 h, 200 nM Na_2_SeO_3_) or rGPX3/rGPX3^U49C^ treatment (D, 24 h, 500 ng/ml). **(E)** Gene expression of *Insr* in 3T3-L1 pre-adipocytes after co-incubation with 500 ng/ml functional (rGPX3) or selenocysteine-to-cysteine mutated enzyme (rGPX3^U49C^) and 1 µM Mithramycin for 24 h. **(F)** GPX activity assay from 3T3-L1 pre-adipocyte lysates after rGPX3/rGPX3^U49C^ treatment (24 h, 500 ng/ml), decrease in NADPH absorption signifies activity. **(G)** Ratio of reduced (GSH) to oxidized (GSSG) glutathione in 3T3-L1 lysates after rGPX3 treatment (24 h, 500 ng/ml). **(H)** Insr mRNA expression in 3T3-L1 cells after heat-inactivated (HI) rGPX3/rGPX3^U49C^ treatment for 24 h. All data are presented as mean ± SEM (N=4 - 8/group). Data in F was analyzed by two-way ANOVA and Šidák’s multiple comparisons test. Significances in A, C, D, and G after 2-tailed Welch’s t-test, *: p< 0.05, ***: p< 0.001.

Our previous data (Hauffe et al., 2020) indicated that the signaling axis leading to an increased InsR expression involves the transcription factor Specificity Protein 1 (SP1). To gain more insights into this regulation, we analyzed the murine *Insr* promoter region and identified a putative SP1 binding site 615 bp upstream of exon 1 (Fig. 4B). To confirm the binding of SP1 to this potential binding site, chromatin immunoprecipitation (ChIP) with an SP1 antibody was performed, coupled with quantitative PCR for this specific promoter region. In line with our previous data, treatment with 200 nM selenite for 24 h increased the enrichment of SP1 to this *Insr* promoter region (Fig. 4C). Importantly, treating cells with either recombinant GPx3 variant did not affect intracellular selenium levels, whereas selenite treatment increased Se by 2.8-fold (Suppl. Fig. S5C). However, treatment with 500 ng/ml of either enzyme is again sufficient to induce a marked 3.5-fold (rGPX3) and 4.6-fold (rGPX3^U49C^) increase in SP1 binding to this promoter region (Fig. 4D), whereas blocking SP1 activity with the GC-box–binding SP1 inhibitor mithramycin A blunted the effect for both enzymes (Fig. 4E). SP1 has been shown to be responsive to changes in the intracellular redox environment, with increased DNA-binding activity observed under conditions of glutathione depletion (Ammendola et al., 1994; Ryu et al., 2003). We therefore investigated whether extracellular GPX3 could, independent of peroxidase activity, alter intracellular redox states and consequently affect SP1 activity.

Thus, we assessed intracellular GPX activity. Interestingly, 24 h rGPX3 or rGPX3^U49C^ treatment both lead to a significant increase in intracellular GPX activity (Fig. 4F). Consistent with GSH consumption during H₂O₂ reduction by intracellular GPX enzymes, rGPX3 treatment decreased the GSH/GSSG ratio by 26 % (Fig. 4G). As rGPX3^U49C^ is functionally inactive, this indicates an increase in intracellular GPX proteins, as opposed to an import of the extracellular enzymes to perform intracellular functions. Furthermore, complete denaturation of the enzymes via heat-inactivation for 10 min at 98 °C (HI-rGPX3 and HI-rGPX3^U49C^) led to loss of function, and, crucially, abrogated the *Insr* upregulation seen with the intact enzymes (Suppl. Fig. S5B, Fig. 4H).

### Extracellular GPX3 enhances insulin signaling and adipocyte function

We next characterized the functional consequences of InsR induction using treatment with extracellular wild-type rGPX3. Probing for phosphorylated InsR and AKT after acute insulin (100 nM) stimulation of 3T3-L1 cells confirmed higher insulin responsiveness after 24 h pre-treatment with rGPX3, as demonstrated by increases in total InsR (+95 %), InsR_Y1150/1151_ (+188 %), and AKT_S473_ (+ 96 %, Fig. 5A). Notably, 24 h of extracellular rGPX3 was also sufficient to significantly increase intracellular GPX1 expression (Fig. 5A), suggesting that extracellular GPX3 might also indirectly influence intracellular redox states. Furthermore, the rGPX3 treatment also led to higher oxygen and glucose consumption in the same pre-adipocyte cell system (Fig. 5B,C). Finally, differentiating 3T3-L1 cells into mature adipocytes in the presence of extracellular rGPX3 showed a 16 % increase in Oil-Red-O accumulation and significantly elevated terminal differentiation markers *Pparγ*, *C/ebpα*, *Ap2*, *Plin1*, and *Glut4*, indicating an increased adipogenic differentiation capacity due to extracellular GPX3 (Fig. 5D,E).

**Figure 5:**
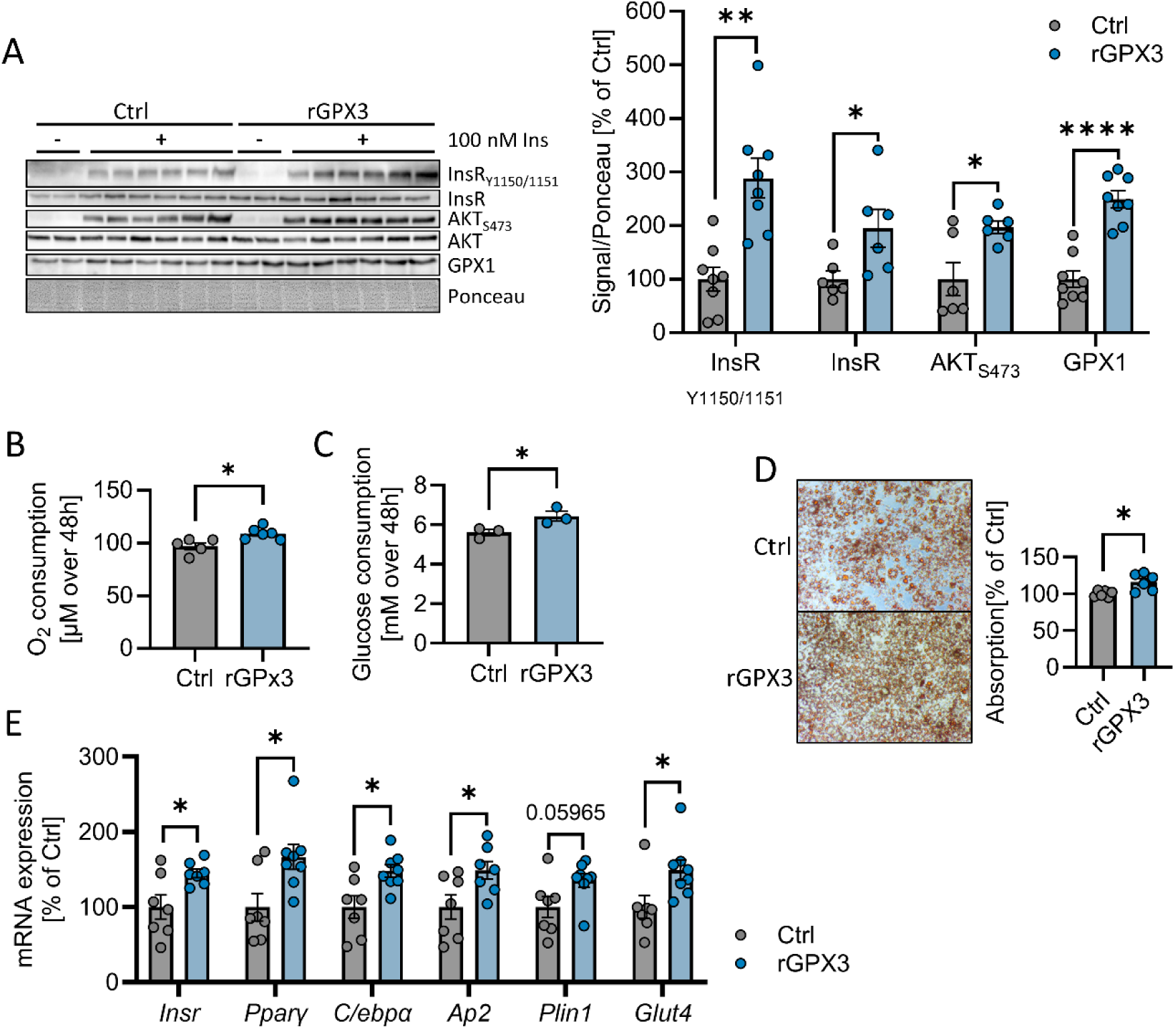
Extracellular recombinant GPX3 improves insulin action and adipocyte functions. **(A)** Phosphorylation, expression and densitometry of InsR, AKT, and GPX1 after acute insulin stimulation of 3T3-L1 pre-adipocytes after rGPX3 treatment (24 h, 500 ng/ml). **(B)** Oxygen consumption of 3T3-L1 pre-adipocytes in the presence of rGPX3 (500 ng/ml) over 48 h. **(C)** Glucose consumption of 3T3-L1 pre-adipocytes in the presence of rGPX3 (500 ng/ml) over 48 h. **(D)** Microscopy and quantification after lysis of Oil-Red-O staining following adipogenic differentiation of 3T3-L1 cells in the presence of 500 ng/ml rGPX3. **(E)** Terminal differentiation marker analysis via qPCR after adipogenic differentiation of 3T3-L1 cells in the presence of 500 ng/ml rGPX3. All data are presented as mean ± SEM (N=6 - 8/group). Significances after 2-tailed Welch’s t-test, *: p< 0.05, **: p< 0.01, ****: p< 0.0001.

### Proof-of-concept validation in mouse-derived primary adipose tissue models

Next, we aimed to extend these findings beyond immortalized cell systems. Accordingly, we performed proof-of-concept experiments in mouse-derived adipose tissue models, using murine *ex vivo*, organoid, and primary cell models.Treating eWAT explants from C57Bl/6N mice with the same 500 ng/ml rGPX3 for 24 h proved sufficient to increase *Insr* expression by 92 %, while heat-inactivation of the enzyme again led to a complete abrogation of *Insr* induction (Fig. 6A). To further validate this observation, we isolated the stromal vascular fraction (SVF) from C57Bl/6N and generated adipocyte organoids (adipoids). Treating those adipoids with 500 ng/ml rGPX3 or rGPX3^U49C^ during the differentiation process resulted in an increase in *Gpx1* gene expression (rGPX3 +39 %, rGPX3^U49C^ + 19 %) which can be, as shown above, indicative of altered intracellular redox activity. Strikingly, either treatment also led to a significant 2.2-fold increase in *Insr* expression (Fig. 6B).

**Figure 6:**
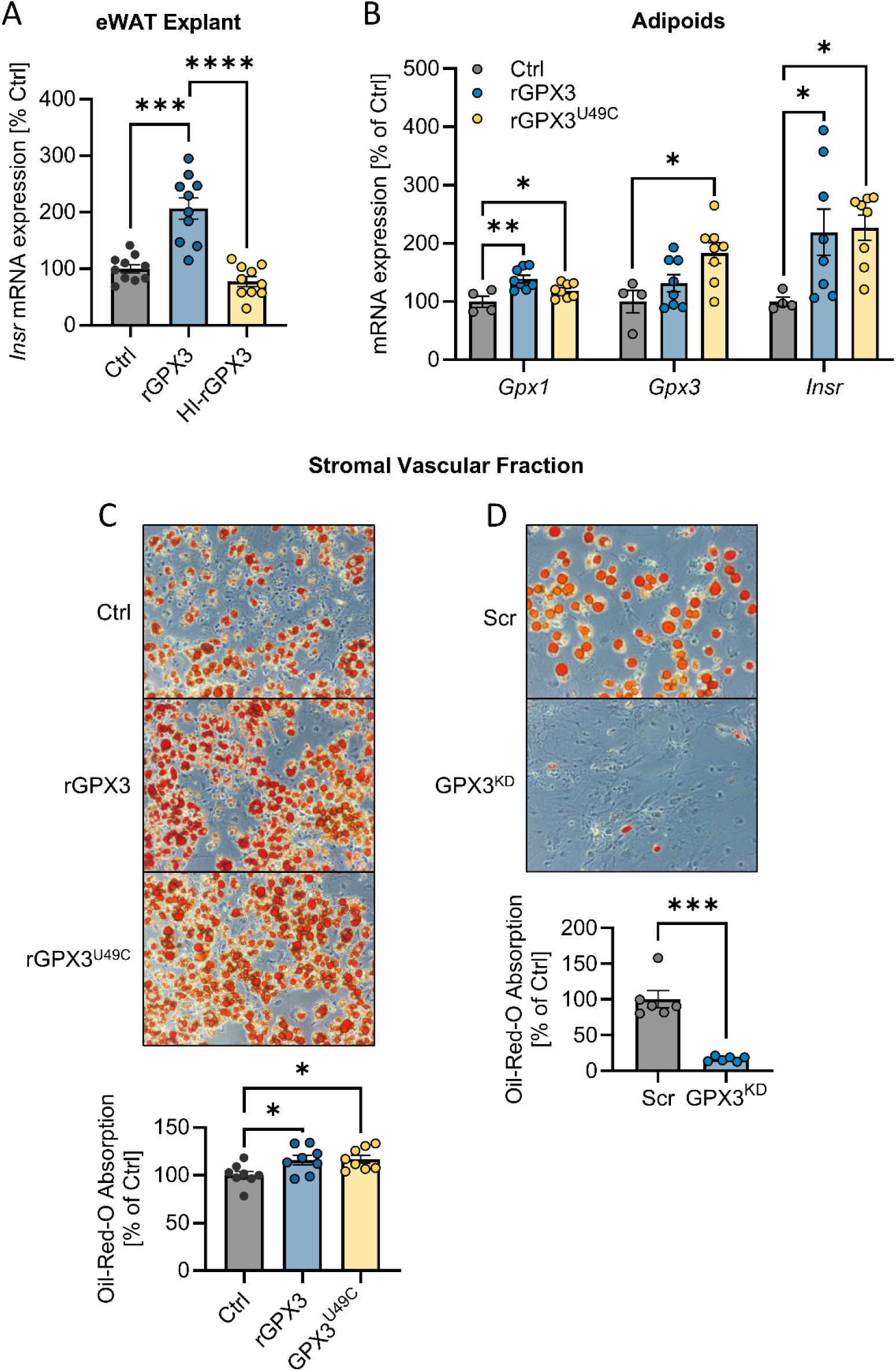
Treatment with extracellular recombinant GPX3 improves adipocyte insulin action in mouse-derived models. **(A)** Gene expression of *Insr* in eWAT explants after rGPX3 or HI-rGPX3 treatment (24 h, 500 ng/ml). **(B)** Gene expression of *Gpx1*, *Gpx3*, and *Insr* in SVF-derived adipose tissue organoids differentiated in presence of rGPX3 or rGPX3^U49C^ (500 ng/ml). **(C)** Microscopy and quantification after lysis of Oil-Red-O staining after adipogenic differentiation of SVF cells pretreated for 48 h before adipogenic induction with 500 ng/ml rGPX3 or rGPX3^U49C^. **(D)** Microscopy and quantification after lysis of Oil-Red-O staining after adipogenic differentiation of SVF cells subjected to scrambled (Scr) or anti-*Gpx3* siRNA (GPX3^KD^) for 48 h before adipogenic induction. All data are presented as mean ± SEM. Significances after 2-tailed Welch’s t-test, *: p< 0.05, **: p< 0.01, ***: p< 0.001, ****: p< 0.0001.

Our 24 h experiments suggest that the InsR induction happens relatively quickly and is present in pre-adipocytes as well as in mature adipocytes. Therefore, we hypothesized that GPX3-associated signaling might pre-dispose adipocyte precursors to more efficient maturation by increasing InsR and, thereby, down-stream signaling. We either exposed SVF cells in 2D culture to 500 ng/ml rGPX3 or rGPX3^U49C^, or depleted endogenous GPX3 by siRNA-mediated GPX3 knockdown only in the proliferation stage before adipogenic induction, not during the maturation process. Interestingly, we observe complimentary phenotypes, as early rGPX3 or rGPX3^U49C^ treatment was sufficient to increase final adipocyte differentiation (measured by Oil-Red-O incorporation after 10 days) by 16 % and 17 %, respectively, whereas early GPX3 knockdown efficiently inhibited adipocyte differentiation (Fig. 6C,D). Collectively, these results identify extracellular GPX3 as a potential paracrine regulator of adipose tissue insulin sensitivity.

## Discussion

In this study, we identified GPX3 as an extracellular regulator of InsR expression and insulin sensitivity in adipose tissue, that seems to act independently of its enzymatic function. By combining human correlation analyses, *in vivo* selenium supplementation, and mechanistic experiments across cell-based and primary adipose models, we provide evidence that GPX3 links adipose tissue redox control to insulin signaling through a transcriptional mechanism involving SP1.

Human adipose tissue datasets reveal a striking association between GPX3 expression and markers of metabolic health, mirroring INSR expression across a broad range of metabolic parameters and increasing dynamically with weight loss. This relationship between metabolic improvement and adipose GPX3 expression is not restricted to dietary or surgical weight loss. Reanalysis of publicly deposited RNA-sequencing data from a recent study of combinatorial incretin-receptor and PPAR agonism in obese mice (Liskiewicz et al., 2026) revealed that eWAT GPX3 expression was increased approximately 1.7-fold by GLP-1:GIP co-agonism (padj = 0.039) and to a comparable extent by GLP-1:GIP:Lanifibranor quintuple agonism (padj = 0.015). Notably, pan-PPAR agonism with lanifibranor alone, administered at a dose equimolar to the conjugate and without effect on body weight in this model, left adipose GPX3 unchanged (padj = 0.98). Pharmacological correction of obesity therefore recapitulates the induction of adipose GPX3 observed after weight loss in humans, and does so in proportion to the degree of metabolic improvement rather than as a direct consequence of PPARγ activation. This seems to contradict the occurrence of PPAR response elements identified in the murine GPX3 promoter (Lee et al., 2008) and suggests that additional, weight-loss-associated signals govern adipose GPX3 expression. Consistent with adipose tissue being a major site of GPX3 synthesis, GPX3 was expressed at approximately 38-fold higher levels in eWAT than in liver in the same dataset. This suggests that GPX3 not only marks healthy adipose tissue, as shown previously (Flehmig et al., 2014), but actively participates in maintaining insulin responsiveness. The contrasting correlations of GPX1 (Suppl. Fig. S1A) reinforce the concept that intracellular and extracellular antioxidant systems exert fundamentally different effects on metabolic signaling (McClung et al., 2004; Huang et al., 2018; Lennicke and Cochemé, 2021).

Prolonged therapeutic selenite supplementation in mice with established obesity improved adipose redox balance, limited adipocyte hypertrophy, and enhanced epididymal fat insulin signaling, ultimately improving glucose tolerance. Notably, these effects were absent in shorter interventions (Hauffe et al., 2022). Attenuated selenium accumulation in obese animals could suggest impaired uptake although a recent meta-analysis found no significant connection between adiposity and Se status (Fontenelle et al., 2022). This suggests, that rather than whole-body Se uptake, it is local dysregulation in tissues that drive Se-related pathologies during metabolic dysfunction, requiring sustained intervention to restore homeostasis.

At the mechanistic level, our data support a model in which extracellular GPX3 enhances InsR expression through SP1 activation in absence of its selenocysteine-dependent activity, suggesting a hormone-like effect. In line, we have published previously that the SP1 inhibitor mithramycin A prevents the *Insr* expression increase after sodium selenite treatment (Hauffe et al., 2020), which is mirrored here for rGPX3 or rGPX3^U49C^ treatment (Fig. 4E). Indeed, underlining the dispensability of selenocysteine, rGPX3 treatment does not increase intracellular selenium levels, suggesting GPX3 does not act as a selenium donor for cells, while denaturation by heat suppressed the effects showing that protein integrity was required (Fig. 4H, Suppl. Fig. S5C). Recombinant GPX3, with or without selenocysteine, at physiological concentrations induced InsR expression across 3T3-L1 cells, primary adipocyte precursors, tissue explants, and adipoids. ChIP experiments confirmed increased SP1 binding to the *Insr* promoter following GPX3 treatment, providing direct evidence for transcriptional regulation. This result suggests that a yet to be identified GPX3-receptor interaction feeds into intracellular redox signaling to influence insulin sensitivity in adipocytes. That a GPX-family protein can retain biological function was recently shown. GPX6, the closest paralog of GPX3, carries selenocysteine in primates but has independently exchanged it for cysteine in at least five placental-mammal lineages. Its physiological function is unknown because the natural Sec-to-Cys substituted form of GPX6 lacks a classical enzymatic GPX activity (Kryukov et al., 2003; Rees et al., 2024). Indeed, modern mouse GPX6 is catalytically indistinguishable from a redox-dead serine variant, yet its fold and glutathione binding are preserved and activity is restored by reintroducing selenocysteine (Rees et al., 2024). Despite this, the cysteine form remains under positive selection and its presence has functional consequences *in vivo*, as it has been shown to interact with mutant huntingtin, which causes Huntington Disease (Shema et al., 2015). GPX enzymes can therefore retain selected-in, non-peroxidase dependent functions, which is consistent with our observation that rGPX3^U49C^ reproduces the InsR, GPX1, SP1 binding, and adipogenesis effects of the wild-type protein.

Despite elevated selenium in multiple tissues, increased GPX3 expression with enhanced insulin signaling was selective to epididymal adipose tissue, highlighting a striking disconnect between selenium supply and functional selenoprotein regulation. These findings are consistent with hierarchical selenoprotein synthesis (Schomburg and Schweizer, 2009), whereby available selenium is preferentially channeled into specific selenoproteins depending on tissue-specific transcriptional and metabolic context. In this framework, subcutaneous adipose tissue, which did show measurable increases in total selenium content, may predominantly incorporate selenium into intracellular selenoproteins such as GPX1 or the selenium transport and storage protein SELENOP, effectively retaining selenium within the tissue. Epididymal adipose tissue, by contrast, showed no increase in total selenium content despite dramatically elevated GPX3 expression and enhanced insulin signaling. This apparent paradox might be explained by the secreted nature of GPX3: selenium incorporated into GPX3 in eWAT would be rapidly released into the extracellular space rather than retained, rendering standard tissue selenium measurements insufficient to detect this functional pool. Indeed, TurboID-based identification showed that adipocyte-derived GPX3 contributes significantly to plasma GPX3 (Plucińska et al., 2026). Thus, the absence of detectable selenium accumulation in eWAT does not reflect a lack of selenium utilization, but rather active and directed incorporation into a secreted effector protein.

This depot-specific response may reflect the distinct metabolic character of epididymal adipose tissue in mice. Compared to subcutaneous depots, eWAT is characterized by higher inflammatory tone, greater metabolic turnover, and stronger association with systemic insulin resistance (Kahn et al., 2019; Börgeson et al., 2022; Kolb, 2022) properties that may prime it to respond preferentially to improvements in systemic redox and nutrient status. In the context of diet-induced obesity and subsequent selenite supplementation, eWAT may therefore function as a metabolic sentinel depot, uniquely positioned to translate subtle changes in selenium availability into adaptive transcriptional responses, including upregulation of the InsR.

Functionally, increased InsR expression translated into enhanced AKT activation in our models, elevated glucose and oxygen consumption, and improved adipogenic differentiation, consistently observed when GPX3 was manipulated within a physiological range and independent of its peroxidase function. However, it is not immediately obvious that a purely transcriptional InsR upregulation model explains all described effects. Studies have shown that receptor tyrosine kinase ligand binding (EGF/PDGF, and by extension insulin via InsR) triggers a NOX-dependent H_2_O_2_ burst that transiently oxidizes PTP1B’s active-site cysteine, which in turn lets the receptor activate. PTP1B is reduced back to its active form by the Glutaredoxin, and Thioredoxin-dependent redox systems acting together, terminating the signal (Coppo et al., 2026; Peng et al., 2016). Thus, it seems plausible that GPX3 signaling might act both via increasing InsR receptor amount, as well as by modulation of intracellular signaling pathways extending individual InsR activity windows.

Our results provide new context for studies reporting both beneficial and detrimental metabolic effects of selenium. Here, GPX3 emerges as a mediator of beneficial effects on insulin sensitivity, whereas excessive GPX1 induction may blunt physiological insulin signaling, potentially explaining selenium’s narrow therapeutic window in human studies (Huang et al., 2018; Morales-Juárez et al., 2026).

Several limitations warrant consideration. Genetic models will be required to definitively establish causality *in vivo*, and conflicting GPX3 whole-body knockout phenotypes (Olson et al., 2009, IMPC MGI:105102) highlight potential compensatory mechanisms. Adipose tissue-specific and inducible GPX3 models will be critical to resolve these issues. Crucially, the precise molecular interaction site of GPX3 on the cell surface also remains to be elucidated.

In conclusion, we identify secreted GPX3 as a paracrine regulator of SP1-dependent InsR expression in adipose tissue independent of its canonic catalytic function. Together with the depot-specific improvement in visceral insulin signaling achieved by restoring adipose GPX3 *in vivo*, these findings suggest GPX3 is a novel paracrine mediator of metabolic health and a potential therapeutic target to reinstate white adipose tissue insulin sensitivity in obesity and type 2 diabetes.

## Methods

### Data analysis from publicly available repositories

Data from adiposetissue.org was extracted from version v1.0.0 in December 2025. For correlations of gene expression and clinical parameters, we used the META sc cohort, traits as shown in the figures, and as correlation method the default Spearman option (Zhong et al., 2025; Armenise et al., 2017; Arner et al., 2012, 2018, 2016; du Plessis et al., 2015; Imbert et al., 2022; Johansson et al., 2012; Keller et al., 2017; Kerr et al., 2020; Krieg et al., 2022; MacLaren et al., 2010; Matualatupauw et al., 2017; Nono Nankam et al., 2020; Petrus et al., 2018; Raulerson et al., 2019; Salcedo-Tacuma et al., 2022; Stančáková et al., 2012; Van Bussel et al., 2017; Vink et al., 2017; Winnier et al., 2015). The same settings were used for comparisons of multiple genes. For the weight loss graphs, we used the default Wilcoxon test to assess significant differences. We used the following cohorts: Bariatric surgery (Kerr,

A. (2020) weight loss sc (Kerr et al., 2020)) was chosen because the study included 5-year follow-up data instead of just 2-year (N_baseline_ = 50, N_2y_ = 49, N_5y_ = 38). Please note that the effect of the other available dataset (Petrus et al 2018), was the same at the 2 year follow up time point. For Diet we followed a similar approach, Vink, RG. (2017) weight loss sc included 2 different intervention diets and a 12-week follow up and Johansson, LE. (2012) weight loss 14 weeks follow up (Johansson et al., 2012; Vink et al., 2017). Again, the other studies show the same effect of *GPX3* increase after weight loss, but stop after 8-week follow-up. Diet cohorts only include BMI as a clinical parameter on adiposetissue.org. In the study by Vink et al., participants followed either a low-calorie diet (LCD; 1250 kcal/day N = 27) for 12 weeks or a very-low-calorie diet (VLCD; 500 kcal/day N = 28) for 5 weeks. In Johansson et al, participants followed a low-calorie diet (LCD; 1200 kcal/day N = 9) for 14 weeks. All data was extracted using the export tables function, and visualized using GraphPad Prism version 10. Data from the International Mouse Phenotyping Consortium (IMPC, Wilson et al., 2026) for heterozygous GPX3 knockout mice phenotypes was extracted in December 2025 from the significant phenotypes table (www.mousephenotype.org/data/genes/MGI:105102). Data was downloaded and visualized using GraphPad Prism version 10. The IMPC makes this dataset publicly available under the Creative Commons Attribution 4.0 International license (CC-BY 4.0).

### Animal experiments

Three-week-old male C57BL/6N mice (Charles River Laboratories, Wilmington, MA, USA) were all continuously kept in a 12 h light/dark cycle at a constant temperature of 22 °C, had free access to water, and were fed ad libitum. The high-fat diets (HFD and selenium-rich HFD (SRHFD)) contained 60 % of calories from fat and were manufactured by ssniff Spezialdiäten GmbH, Soest, Germany). The selenium contents are as ∼230 ng Se/g diet (HFD) and ∼670 ng Se/g diet (SRHFD), see Hauffe et al., 2020 for detailed analysis of trace element content. Body weight was monitored weekly. At 4 weeks of age, mice were fed the regular HFD for 8 weeks to induce overweight and insulin resistance. Then, mice were randomly sorted to receive either the same HFD or the SRHFD for further 20 weeks. Insulin tolerance test was performed 14 weeks after diet switch to HFD or SRHFD by intraperitoneal injection of insulin (0.75 U/kg bodyweight). The oral glucose tolerance test was done 16 weeks after diet switch, animals were starved for 16 h overnight and 2 g glucose/kg bodyweight was administered by gavage. Peak glucose excursion was defined as the maximum recorded value across all time points for a given animal, expressed as the absolute difference above the individual baseline value at t = 0 min (Peak above baseline = Peak value − Baseline). The rate of change during the ascending phase (ROC up) was calculated as the slope from baseline to peak, defined as (Peak value − Baseline) / (Time to peak − 0), yielding units of mM per minute. All per-animal values were calculated using that animal’s own baseline. For final time points, 6 mice per diet group were chosen randomly for insulin stimulation experiments. at At 32 weeks of age (20 weeks after diet switch) the remaining 14 mice were killed by cervical dislocation, tissues were harvested, weighed, and processed for histological, biochemical, and molecular analyses.

Vena cava insulin injection to assess local insulin sensitivity was performed at 32 weeks of age (20 weeks after diet switch). Mice were starved overnight and anaesthetized the following morning. Then, 5 U of insulin or saline as control were injected directly into the vena cava. Organs were harvested after the time points quoted in parentheses: Liver (2 min), quadriceps (5 min), gonadal and subcutaneous white adipose tissue (5 min), and brain and brown adipose tissue (10 min).

### Declaration of Approval

All animal procedures were conducted according to German laws for the protection of animals and in compliance with protocols approved by local governmental authorities (State Agency of Environment, Health and Consumer Protection, State of Brandenburg, Germany, approval number 2347-30-2020). *Adipocyte area analysis*

Fresh adipose tissue samples were submerged in 4 % paraformaldehyde for 24 h at 4°C, washed in PBS and embedded in paraffin. Slices were fixed on glass slides and stained with hematoxylin and eosin. Microscopy pictures were taken with a 10x magnification objective on a microscope (Zeiss Axio Vert. A1). Adipocyte size and morphology were quantified using a custom image analysis tool (AdiQuant; available at GitHub, link provided in Data and Resource Availability). The software applies a multi-step image processing pipeline to histological images. Briefly, images were converted to grayscale and contrast was enhanced using Contrast Limited Adaptive Histogram Equalization (CLAHE; clip limit 2.0, tile grid 8×8), followed by Gaussian filtering (5×5 kernel) to reduce noise. Adipocytes were segmented using adaptive Gaussian thresholding based on local pixel neighborhoods (31×31 pixels). Segmentation masks were refined using morphological opening and closing operations (3×3 kernel). Touching adipocytes were separated using Euclidean distance transformation and watershed segmentation. Individual cells were quantified using region-based analysis to determine adipocyte area and perimeter. Areas were calculated from pixel counts and converted to µm² using calibrated scale parameters. To exclude artifacts, segmented objects smaller than 100 pixels were removed prior to analysis.

### Generation of recombinant enzymes: Glutathione Peroxidase 3 (rGPX3) and selenocysteine-deficient GPX3 (rGPX3^U49C^)

Recombinant mouse GPX3 (UniProtKB – P46412, 226 aa) without signal peptide (1-24 aa) was cloned after codon optimization for E. coli expression, using a gene cassette insert ordered from Integrated DNA Technologies, Inc. (IDT), with the synthetic gene transferred into the pABC2 plasmid developed previously by us (Cheng and Arnér, 2017), to generate fusion proteins linked with an N-terminal His-tagged Sumo protein. For recombinant GPX3 without selenocysteine, site specific mutagenesis on the pABC2a-MmGPX3 plasmid using NEBaseChanger methodology (TAG → TGT) was used to exchange selenocysteine at position 49 with a cysteine, yielding pABC2a-MmGPX3 U49C. See supplementary table S1 for sequence details. The final expression plasmid was transformed into E.coli C321.ΔA strain (Lajoie et al., 2013).

The recombinant GPX3 was expressed in E. coli C321.ΔA. Expression conditions and purification of target protein from bacterial cell lysates using affinity chromatography were basically as described previously (Cheng and Arnér, 2017). Briefly, 40 ml aliquot of overnight culture of the transformed bacteria was used to inoculate 2 L terrific broth (TB) media supplemented with 50 µg/ml streptomycin in 5 L Erlenmeyer flasks. The C321.ΔA cells were propagated at 33°C in shaking incubators. Around 5-6 h after inoculation, when the optical density of the cultures reached 1.2-1.4 at 600 nm, the temperature was decreased to 24°C and Na2SeO3 was added at 5 µM final concentration from 20 mM stock solution. The protein expression was initiated by adding 0.5 mM IPTG and the culturing was continued overnight. After propagation, bacterial cells were harvested by centrifugation and suspended in binding buffer for nickel-column affinity chromatography (50 mM Tris-HCl, 250 mM NaCl, 25 mM imidazole, pH 7.5). Extraction of proteins was achieved by sonification, and the cell lysate was centrifuged. The supernatant containing the soluble fraction of proteins was loaded to HisTrapTM HP 5 ml column (cytiva) installed on an ÄKTA explorer FPLC system (Cytiva Life Sciences). After washing out the contaminating proteins from the column, the 6xHis-SUMO-GPX3 was eluted from the column under isocratic condition using 40 % elution buffer (50 mM Tris-HCl, 250 mM NaCl, 500 mM imidazole, pH 7.5). Subsequently the fusion tag was cleaved from the N-terminal part of the protein by 6xHis-ULP1 SUMO protease made in-house. The resulted mixture of the target GPX3, 6xHis-ULP1 and 6xHis-SUMO tag was re-loaded to the same HisTrapTM HP 5 ml column to bind the His-tagged, contaminating proteins to the column, and the purified non-tagged GPX3 could be retrieved in the flow-through. After buffer change and concentration, the recombinant GPX3 was stored in 25 mM Tris-HCl (pH 8.0) with 250 mM KCl, 2 mM 2-Mercaptoethanol and 10 % glycerol. The purity of the isolated proteins was over 95 % as judged by SDS-PAGE analysis.

### Cell culture

White 3T3-L1 pre-adipocyte cells were cultured in Gibco DMEM GlutaMAX™ (Thermo Fisher Scientific Inc., USA) containing 450 mg/dL glucose, and 10 % fetal bovine serum (FBS), 1 % penicillin/streptomycin (Thermo Fisher Scientific Inc., USA), and 1 % pyruvate (Thermo Fisher Scientific Inc., USA) at 37 °C and 5 % CO_2_. To assess acute insulin sensitivity cells were stimulated with 100 nM insulin for 5 min after 3 h FBS deprivation. 3T3-L1 differentiation into mature adipocytes was done by culturing until confluency. 24 h after confluency, differentiation was induced by addition of 0.5 mM 3-Isobutyl-1-methylxanthin (IBMX, Merck KGaA, Germany), 1 µM Dexamethasone (Merck KGaA, Germany), 10 µg/mL insulin (Merck KGaA, Germany), and 1 µM Rosiglitazone (Cayman Chemical, USA) for 48 h. Cells were then cultured in full media in presence of 10 µg/mL insulin for further 8 days. Stimulation with selenocysteine-containing or selenocysteine-deficient recombinant GPX3 (rGPX3 or rGPX3^U49C^) was done for indicated time-points at a concentration of 500 ng/ml. An equal volume of rGPX3 reconstitution buffer was used as control (see below for composition). Assuming every rGPX3 polypeptide incorporated selenocysteine at the UAG codon, this would equal 22.3 nM selenium supplementation. To inhibit SP1 activity, 1 µM mithramycin A (Merck KGaA, Germany), was used in combination with other treatments for the same durations.

### Oxygen and glucose consumption

Oxygen consumption was continuously monitored using the SensorDish® Reader (PreSens, Germany), which enables non-invasive, time-resolved measurements of dissolved oxygen in cell culture plates. Oxygen consumption over 48 h was determined by: -1 * (O_2_^48h^ – O_2_^0h^). Within the same time-frame, media glucose was measured using a commercial glucometer (Contour Glucometer XT, Bayer, Germany) and glucose consumption over 48 h was determined by: -1 * (Glucose^48h^ – Glucose^0h^).

### Total Glutathione Peroxidase Activity Assay

Total glutathione peroxidase (GPX) activity was measured in tissue and cell lysates using a coupled enzymatic assay based on NADPH oxidation as described previously (Florian et al., 2010). In brief, samples or homogenization buffer (blank) were incubated with assay buffer containing Tris/HCl, EDTA, sodium azide, Triton X-100, reduced glutathione, NADPH, and glutathione reductase at 37 °C. The reaction was initiated by addition of hydrogen peroxide, and the decrease in NADPH absorbance was monitored kinetically at 340 nm using a plate reader. GPX activity was calculated from the rate of NADPH consumption and normalized to total protein concentration of each sample.

### Determination of GSH and GSSG

Determination of reduced glutathione (GSH) and oxidized glutathione disulfide (GSSG) was done with the GSH/GSSG-Glo™ Assay (Promega Corporation, USA), according to manufacturer’s instructions.

### WAT explants and Stromal vascular fraction (SVF) - isolation and differentiation

For explant culture, epidydimal WAT from 10-16 weeks old male C57Bl/6N mice was dissected, cut into ∼30 mg pieces and briefly washed in PBS. Explant pieces were then deposited in individual wells of a 24 well plate in Gibco DMEM GlutaMAX™ and stimulated with 500 ng/µL rGPX3, rGPX3^U49C^ or equal volume of resuspension buffer (see above) for 24 h. From the same mice, SVF was isolated from subcutaneous white adipose tissue depots by collagenase II digestion (1 mg/ml in 3.5 % BSA, 45 min at 37 °C), followed by pelleting of the pre-adipocytes by centrifugation. SVF-derived pre-adipocytes were differentiated into white adipocytes using an optimized 2D monolayer protocol adapted from (Scott et al., 2024). Cells were seeded in growth medium consisting of a 7:3 mixture of high-glucose DMEM and DMEM/F12 supplemented with 3% FBS and 1 % penicillin/streptomycin, at a density allowing cells to reach approximately 90 % confluence within 1-2 days. Additionally, cells were supplemented with either 500 ng/ml rGPX3 or buffer control during the expansion time. Differentiation was initiated at near-confluence without a prolonged post-confluent arrest. Importantly, rGPX3 treatment was stopped after the expansion time and not resumed during the rest of the differentiation. From day 0 to day 3, cells were cultured in induction medium (base medium as above) supplemented with 0.5 mM IBMX, 1 µM dexamethasone, 100 nM human insulin, 5 µM rosiglitazone, 33 µM biotin (Merck KGaA, Germany), and 17 µM pantothenate (Merck KGaA, Germany). After 72 h, medium was replaced with maintenance medium containing base medium supplemented with 100 nM insulin, 33 µM biotin, and 17 µM pantothenate during days 3–5. Maintenance medium was refreshed every 48 h until day 14. Adipose tissue organoids (Adipoid) culture was performed as described elsewhere (Taylor et al., 2020).

### Knockdown of GPX3 via siRNA

Knockdown experiments were performed with smartPOOL siRNA (GPX3: M-042578-01, nontarget: D-001810-01-05, Horizon Discovery Ltd., USA) according to the manufacturer’s guidelines. SVF cells were treated with the respective siRNAs only during the proliferation stage, before adipogenic induction. *Oil-Red-O staining*

Adipocyte differentiation and maturation were monitored by phase-contrast microscopy, and lipid accumulation was assessed at day 10 using Oil-Red O staining. Cells were fixed in 4 % Paraformaldehyde for 20 min, washed with PBS, and stained with Oil Red O (0.3 % in 60 % isopropanol/40 % H_2_O) for 30 min. A representative microscopy picture was taken. Subsequently, cells were lysed in 100 % isopropanol to solubilize Oil Red O dye. The absorbance of the resulting solution was measured in a microplate reader at 520 nm.

### RNA isolation, cDNA synthesis, and quantitative real-time PCR

RNA was isolated using NucleoZOL compound (Macherey-Nagel, GmbH & Co. KG, Germany), following the manufacturer’s instructions. Reverse transcription of RNA to cDNA was performed with the Biozym Scientific GmbH cDNA synthesis kit, according to instructions. qPCR was done with Biozym Blue S’Green qPCR Kit on a C1000 Touch Thermal Cycler with CFX384 Real-Time system (Bio-Rad Laboratories GmbH). Beta-Actin was used as a housekeeping gene and data analyzed using a custom software utilizing the ddCt method (CtQuant; available at GitHub, link provided in Data and Resource Availability). Primer sequences can be found in supplementary material, table S2.

### Protein isolation, SDS-PAGE and Western Blotting

Proteins from cells or tissues (ground in liquid nitrogen using mortar and pestle) were isolated by cell/tissue lysis in RIPA buffer, supplemented with protease and phosphatase inhibitors (cOmplete™ Protease Inhibitor Cocktail, and PhosSTOP™, both Merck KGaA, Germany), and subsequent centrifugation. For SDS-PAGE, 15 µg Protein was loaded per gel in Laemmli loading buffer, and separated on a 12 % acrylamide gel. Western blotting was performed by tank blotting. Following SDS-PAGE, proteins were transferred to a membrane using a tank blotting system. Prior to transfer, membranes were activated in methanol for 30–60 s, briefly rinsed with distilled water, and equilibrated in transfer buffer (48 mM Tris, 39 mM glycine, 0.08 % SDS, 20 % methanol). Gels, membranes, and filter papers were pre-equilibrated in transfer buffer before assembly of the transfer sandwich. Air bubbles were removed by gently rolling the sandwich. Blotting was performed at fixed voltage of 90 V for 3 h. Please note, due to technical issues with the tank blotting system, we used a semi-dry blotting system for blots shown in supplementary figure S3. In this case, proteins were transferred at 1.5 mA/cm² for 2 h in transfer buffer (25 mM Tris, 100 mM glycine, 0.08 % SDS, 20 % methanol). After transfer, membranes were blocked in 5 % milk prepared in TBST (0.05 % Tween-20 in TBS) for 30–60 min at room temperature. After washing with TBST, membranes were incubated with primary antibodies diluted in TBST containing 0.2 % milk overnight at 4 °C with gentle agitation. All primary antibodies were diluted at 1:1000, secondary antibodies 1:2000. Antibodies used in this manuscript can be found, with the corresponding Research Resource Identifier (RRID), in the supplementary material, table S2. Carbonylated proteins were detected by derivatization with 20 mM 2,4-dinitrophenylhydrazine (DNPH) in 2 M HCl and subsequent binding to anti-DNP antibody.

Ponceau staining was used as loading control and normalization factor. Western blot band intensities were quantified using a custom image analysis tool (BlotQuant; available at GitHub, link provided in Data and Resource Availability). Images were converted to grayscale and, if necessary, rotationally aligned using bicubic interpolation to ensure vertical lane orientation. Regions of interest (ROIs) encompassing bands of interest were manually defined and subdivided into individual lanes based on replicate number, with optional manual adjustment to account for irregular spacing. Band intensities were quantified using integrated density measurements after local background subtraction, determined by the median intensity within each lane ROI. Signal was defined as pixels exceeding a dynamic threshold (background + 0.2 × standard deviation), allowing exclusion of background noise. Target protein signals were normalized to corresponding loading control intensities from the same lane prior to statistical evaluation.

### Chromatin immunoprecipitation (ChIP)

Identification of putative SP1 binding site in the murine *Insr* promoter (assembly GRCm39) was done by searching for core GC box elements up to 1500 base-pairs upstream of *Insr* exon 1, using SnapGene (version 8.2.2). ChIP was performed as described in (Schupp et al., 2009): Adherent cells at 70–80 % confluence were cross-linked directly in culture medium with 1 % formaldehyde (270 µL of 37 % formaldehyde per 10 mL medium) for 10 min at room temperature with gentle agitation. Cross-linking was quenched by adding glycine to a final concentration of 125 mM and incubating for 5 min. Cells were washed twice with ice-cold PBS, scraped on ice, collected in 15 mL tubes, and centrifuged at 1,350 × g for 5 min at 4°C. Cell pellets were stored at −80°C until further processing. Cell pellets were resuspended in hypotonic lysis buffer and homogenized using a B-type pestle (≥20 strokes). Nuclei were pelleted by centrifugation at 3,000 × g for 5 min at 4°C, washed twice with hypotonic buffer, and lysed in SDS lysis buffer (5× pellet volume) supplemented with protease inhibitors. After incubation on ice for 20 min, chromatin was sheared by sonication (2 cycles of 30 s on/30 s off on ice, amplitude 100). Lysates were cleared by centrifugation at 3,000 × g for 5 min at 4°C, and supernatants were used for ChIP.

DNA concentration was determined by Nanodrop (OD260), and 100–400 µg DNA per sample was diluted to 1 mL in ChIP dilution buffer A containing protease inhibitors. Samples were incubated overnight at 4°C with the indicated antibodies on a rotating platform. Protein A/G magnetic beads (25 µL) were then added and incubated for 2 h at 4°C. Beads were sequentially washed with ChIP dilution buffer B, high-salt buffer (500 mM NaCl), LiCl buffer, and TE buffer, with each wash performed for 5 min at room temperature with rotation.

Chromatin complexes were eluted in 250 µL elution buffer and cross-links were reversed by overnight incubation at 65°C. Eluates were supplemented with 250 µL TE buffer, Proteinase K (final concentration 0.3 mg/mL), and glycogen (20 µg), followed by incubation at 37°C for 2 h. After addition of NaCl (final concentration 200 mM), DNA was purified by phenol-chloroform-isoamyl alcohol extraction and ethanol precipitation. Pellets were washed with 75% ethanol, air-dried, and resuspended in 30–50 µL RNase-free water. RNA was removed by RNase A treatment (500 ng/mL, 37°C, 45–60 min).

Enrichment of genomic sites in input and SP1 ChIP was determined by qPCR, normalized to a site near the Insulin (*Ins)* gene, and evaluated using standard curves for each primer pair (Suppl. table S4).

### Inductively Coupled Plasma – Mass Spectrometry (ICP-MS) for trace element analysis

Tissue samples of mice, fed with either HFD or SRHFD feed for 20 weeks after established obesity, were weighed into TFA vessels and subjected to acid-assisted microwave digestion (Mars 6, CEM, Kamp-Lintfort, Germany) using 65 % HNO3 (Suprapur, Merck, Billerica, MA, USA) and 30 % H2O2 (Merck, Billerica, MA, USA), supplemented with germanium, 77selenium (final concentration: 10 µg/L) and rhodium (final concentration: 1 µg/L) as internal standards. Digestion was carried out with the following parameters: 650 W, 15 min ramp, 20 min at 200 °C, yielding a clear solution. Cell pellets were subjected to the same digestion protocol. Cell numbers were determined using the DeNovix CellDrop FLi system prior to pelleting to allow normalization of metal contents. Selenium, zinc, iron, manganese, and copper were quantified using an ICP-MS/MS (8800 ICP-QQQ-MS, Agilent Technologies, Waldbronn, Germany). Experimental conditions for ICP-MS/MS measurements are listed in Supplementary Tables S5 and S6. To validate digestion and metal quantification, certified reference materials fish muscle (ERM-BB 422) and pig kidney (ERM-BB 186) were subjected to the same protocol (recoveries listed in Supplementary Table S5). Selenium was quantified via isotope dilution analysis as described previously (Rodríguez-González et al., 2005). The instrument was tuned daily for maximum sensitivity on 80→96Se and a doubly charged ratio <2 % (140Ce2+/140Ce+) as well as a background of <0.1 CPS.

Next, all measured trace element concentrations were normalized to tissue-specific HFD control means (% HFD control). Normalized tissue values were averaged per animal to generate a single composite trace element value, ensuring that statistical analyses were performed using the mouse as the experimental unit.

### Signal peptide prediction

To determine the presence of a secretion signal peptide, we utilized SignalP 5.0 (https://services.healthtech.dtu.dk/services/SignalP-5.0/) in December 2025 as described elsewhere (Almagro Armenteros et al., 2019). We used the murine amino acid sequence of GPX3 (uniprot.org ID P46412) as input.

### Protein structure prediction

The structure predictions of proteins was done in December 2025 on Alphafold 3 (https://alphafold.ebi.ac.uk/, Fleming et al., 2025), using amino acid sequences of GPX3 (uniprot.org ID P46412) and GPX1 (uniprot.org ID P11352), respectively.

### Statistical analysis

Data are presented as mean ± SEM unless otherwise indicated. Statistical significance was determined using appropriate tests based on experimental design (e.g., Welch’s t-test, ANOVA with post hoc correction). A p-value < 0.05 was considered statistically significant. All data analysis from adiposetissue.org was done through the implementation of statistical tests within the portal. Details for the tests of datasets used here are written in the respective figure legends.

### Data and Resource Availability

All own data generated or analyzed during this study are included in the published article (and its online supplementary files). Raw gene-level counts from GSE314029 (Liskiewicz et al., 2026) that are referenced in the discussion were retrieved from GEO and subset to the epididymal white adipose tissue samples. Counts were normalised by the median-of-ratios method, and each treatment group was compared with vehicle using negative-binomial Wald tests with Benjamini–Hochberg correction (pyDESeq2 v0.5.4). Reported values are the GPX3 log2 fold-change and adjusted p-value per contrast. Data from adiposetissue.org is available at their domain. The adipocyte analysis software AdiQuant, the Western blot quantification software BlotQuant, the qPCR analysis tool CtQuant, and the ITT/GTT analysis tool TolQuant, including standalone executables, source code, documentation, and usage instructions, are available at the respective GitHub repositories and archived via Zenodo (AdiQuant: github.com/RobHauffe/AdiQuant, DOI: 10.5281/zenodo.18657077, BlotQuant: github.com/RobHauffe/BlotQuant, DOI: 10.5281/zenodo.19691450, CtQuant: github.com/RobHauffe/CtQuant, DOI: 10.5281/zenodo.22869508, TolQuant: github.com/RobHauffe/TolQuant, 10.5281/zenodo.22869542). These tools are free for academic and non-commercial research use.

## Declaration of interests

The authors have no competing interests to declare.

## Author contributions

R.H.: Conceptualization, Data curation, Formal analysis, Funding acquisition, Investigation, Methodology, Software, Supervision, Validation, Visualization, Writing – original draft, Writing – review & editing. M.G.: Investigation, Methodology, Writing – review & editing. S.C.: Investigation, Methodology, Writing – review & editing. J.S.M.: Investigation, Methodology, Writing – review & editing. C.S.: Investigation, Methodology, Writing – review & editing. M.M.: Investigation, Methodology, Writing – review & editing. V.K.: Investigation, Methodology, Writing – review & editing. A.A.: Investigation, Methodology, Writing – review & editing. E.A.: Funding acquisition, Project administration, Resources, Supervision, Writing – review & editing. A.K.: Conceptualization, Funding acquisition, Project administration, Resources, Supervision, Writing – original draft, Writing – review & editing.

## Declaration of generative AI and AI-assisted technologies in the manuscript preparation process

During the preparation of this work, the authors used Anthropic’s Claude Opus 5 Model in order to search and synthesize complex literature, improve language and readability of the methods sections, and assist in coding the analysis tools published in this article. After using this tool, the authors reviewed and edited the content as needed and take full responsibility for the content of the published article. The validity of the analysis toll outputs was verified against previously used standard analysis methods.

## Acknowledgments

We would like to thank Lydia Häußler for technical assistance and Dr. Michael Schupp for invaluable help setting up the Chromatin-immunoprecipitation protocols.

## Funding

This work was supported by two Deutsche Diabetes Gesellschaft grant (to R.H. and A.K., respectively), and Deutsche Forschungsgemeinschaft (DFG) grant project KL 2399/8-1 (to A.K.) and MA 9681/1-1 (to M.M.). The production and work with recombinant GPX3 variants were supported by Karolinska Institutet, The Swedish Cancer Society (24 3482 Pj), and The Hungarian National Research, Development and Innovation Office (NKFIH), under the National Laboratories Program (National Tumor Biology Laboratory (2022-2.1.1-NL-2022-00010)), the Hungarian Thematic Excellence Program (under project TKP2021-EGA-44) and Project Grant K 146277 (to E.S.J.A).

## Notes

### Competing Interest Statement

The authors have declared no competing interest.

