## SupplementaryFiguresandMaterial for "Extracellular GPX3 drives insulin receptor expression in adipose tissue independent of its peroxidase activity"

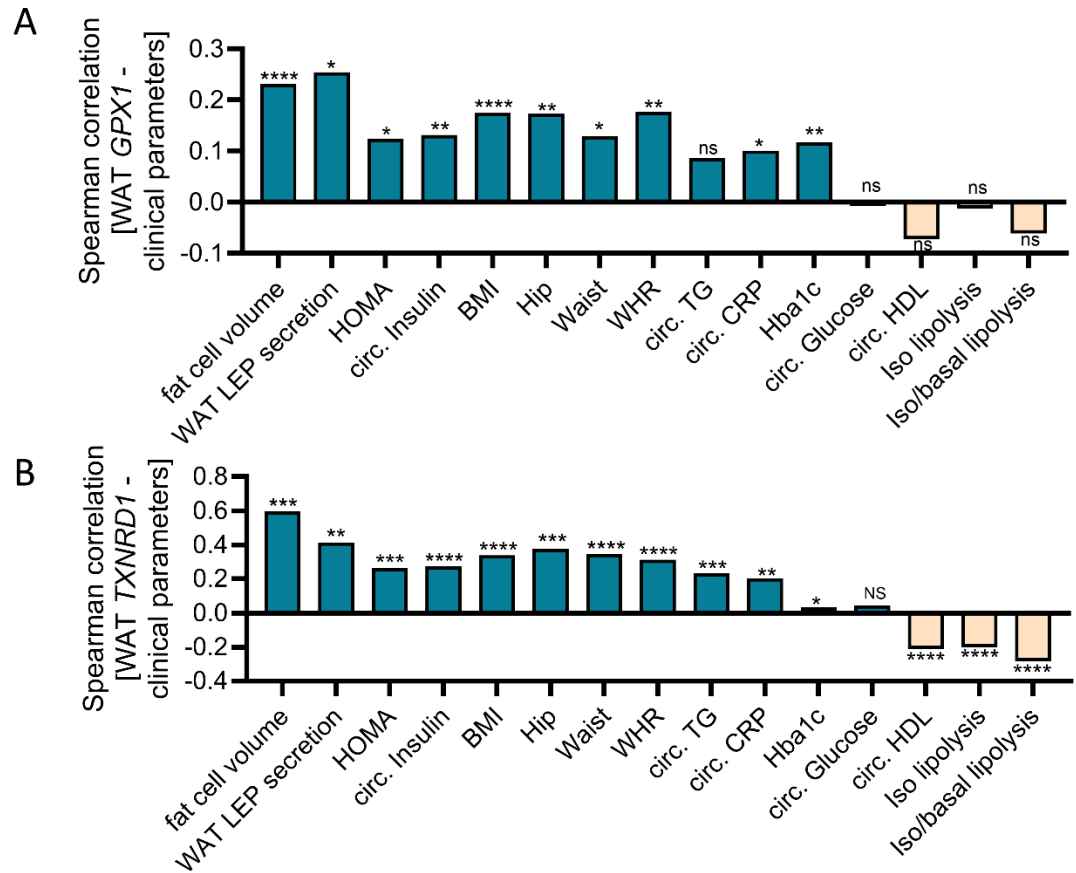

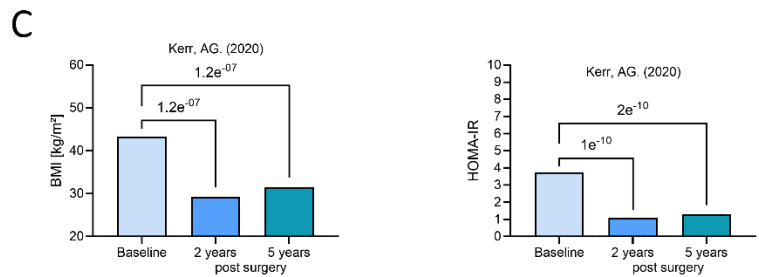

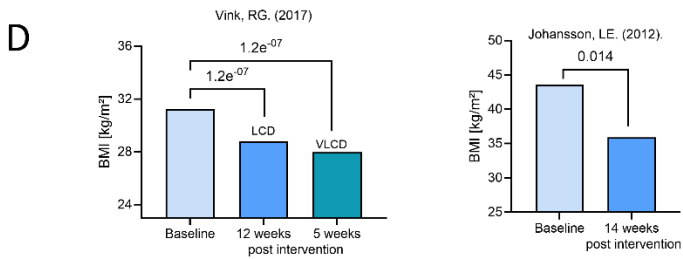

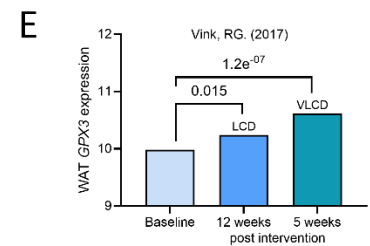

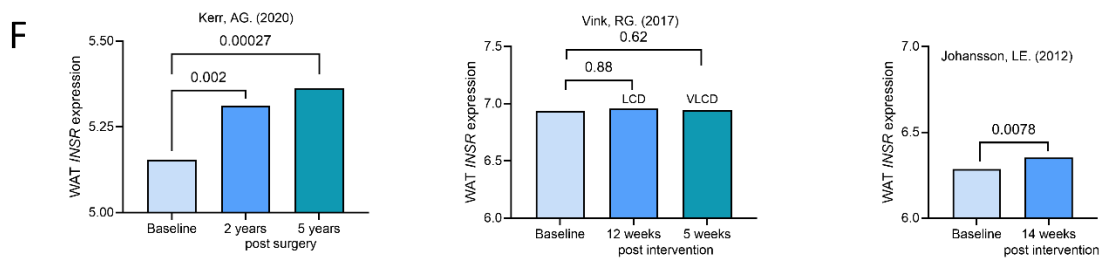

**Supplementary Figure S1:** Association of *GPX1*, *GPX3*, *TXNRD1*, or *INSR* and metabolic traits, weight outcomes of weight-loss interventions: **(A,B)** Correlation of WAT *GPX1* (A) or *TXNRD1* (B) expression and metabolic traits. Bar height represents Spearman correlation coefficient ($\rho$). Asterisks indicate $-\log_{10}(p)$ category (\* $p \geq 10$-20, \*\* $p \geq 20$-30, \*\*\* $p \geq 30$-60, \*\*\*\* $p > 60$). **(C)** BMI and HOMA-IR decreases after weight loss through bariatric surgery **(D)** BMI decreases after weight loss through diet. **(E)** *GPX3* expression associates with weight loss through diet **(F)** *INSR* expression associates with weight loss through bariatric surgery or diet. Statistical differences between baseline and follow-up time points were assessed using the Wilcoxon signed-rank test, as implemented in adiposetissue.org. All data extracted from adiposetissue.org v1.0.0. See methods section for references.

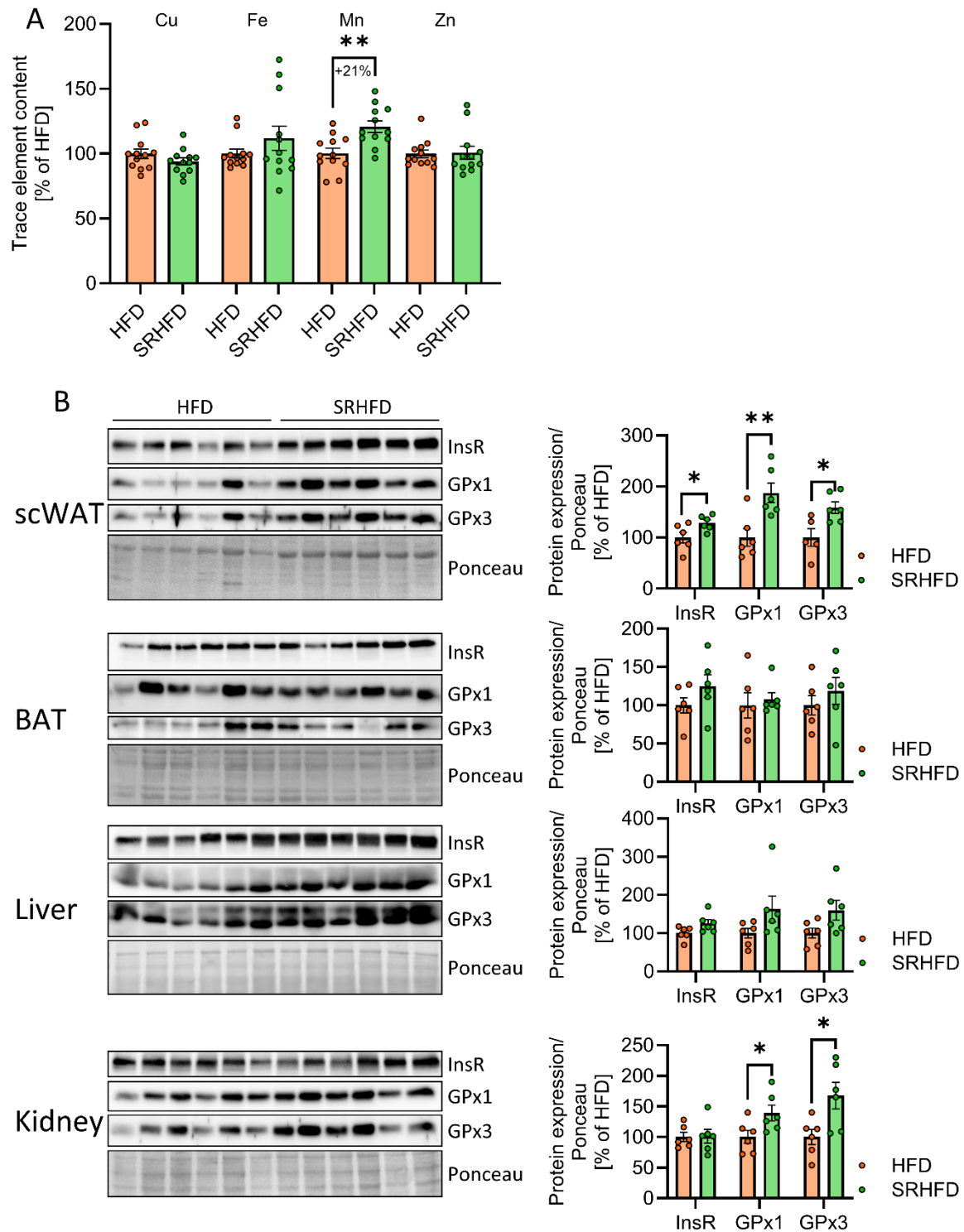

**Supplementary Figure S2:** InsR and GPX expression in various tissues. Data in A and B compares male C57BL/6N mice fed either HFD or SRHFD after established obesity for 20 additional weeks. **(A)** Total trace element contents of copper (Cu), iron (Fe), manganese (Mn) and zinc (Zn), data aggregated from liver, kidney, eWAT, scWAT, and BAT trace element levels per mouse. **(B)** Protein expression and

densitometric analysis of InsR, GPX1 and GPX3 in scWAT, BAT, liver, and kidney lysates. All data are presented as mean ± SEM. Significances after 2-tailed Welch's t-test, \*: $p < 0.05$, \*\*: $p < 0.01$.

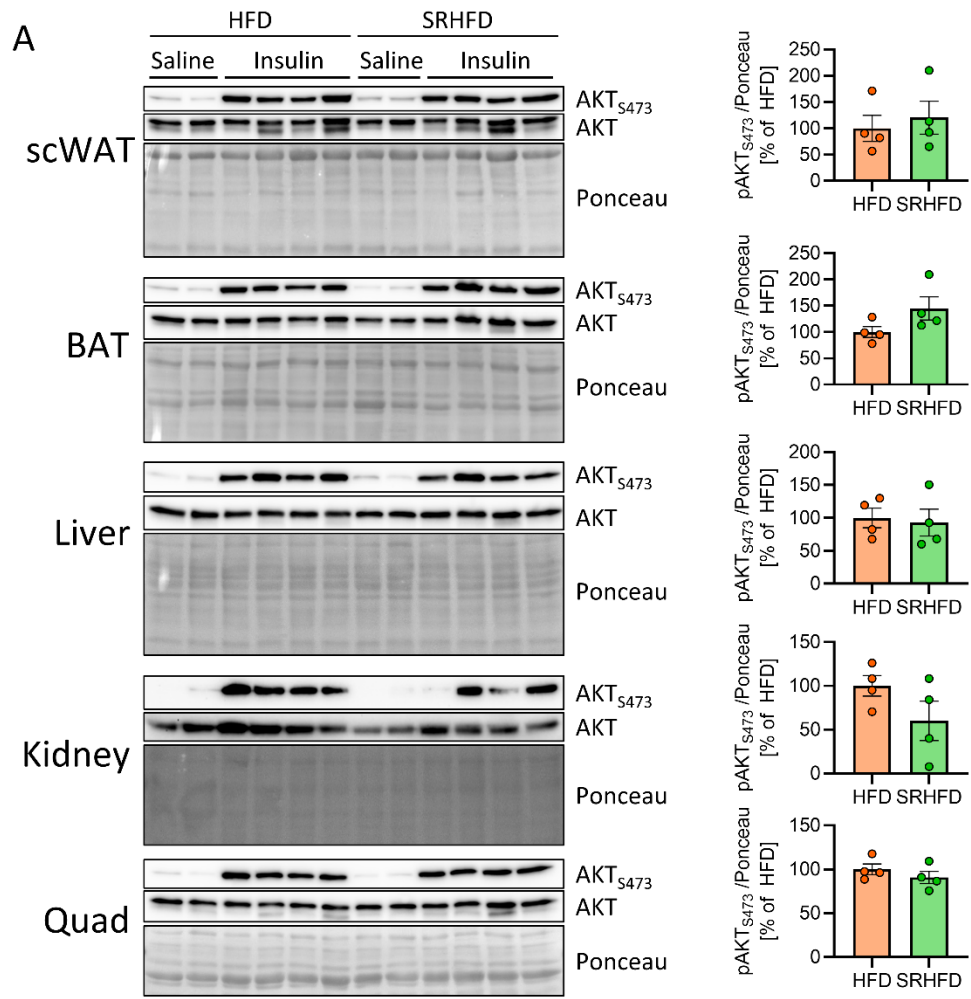

**B**

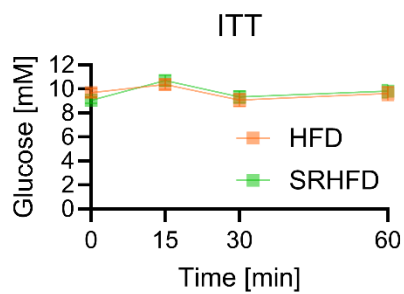

**C**

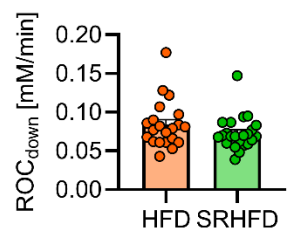

**D**

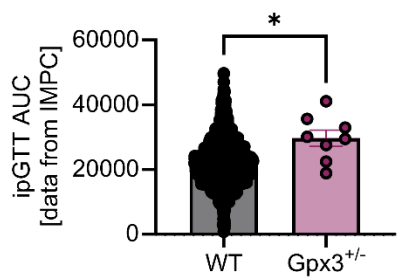

**Supplementary Figure S3:** Insulin sensitizing effect is restricted to visceral WAT. **(A)** Protein phosphorylation, expression and densitometric analysis of AKT in scWAT, BAT, liver, kidney, and quadriceps lysates after vena cava injection of saline or insulin in male C57BL/6N mice fed either HFD or SRHFD after established obesity for 20 additional weeks. **(B)** Blood glucose levels during an insulin tolerance test after 14 weeks of HFD or SRHFD. **(C)** Rate of change from peak to last time point ($ROC_{down}$) during the oGTT shown in Fig. 3E. **(D)** Area under the curve from an ipGTT of wildtype (WT) and heterozygous, whole-body GPX3 knockout mice (data from IMPC, http://www.mousephenotype.org/data/genes/MGI:105102). All data are presented as mean ± SEM. Significances after 2-tailed Welch's t-test, \*: $p < 0.05$.

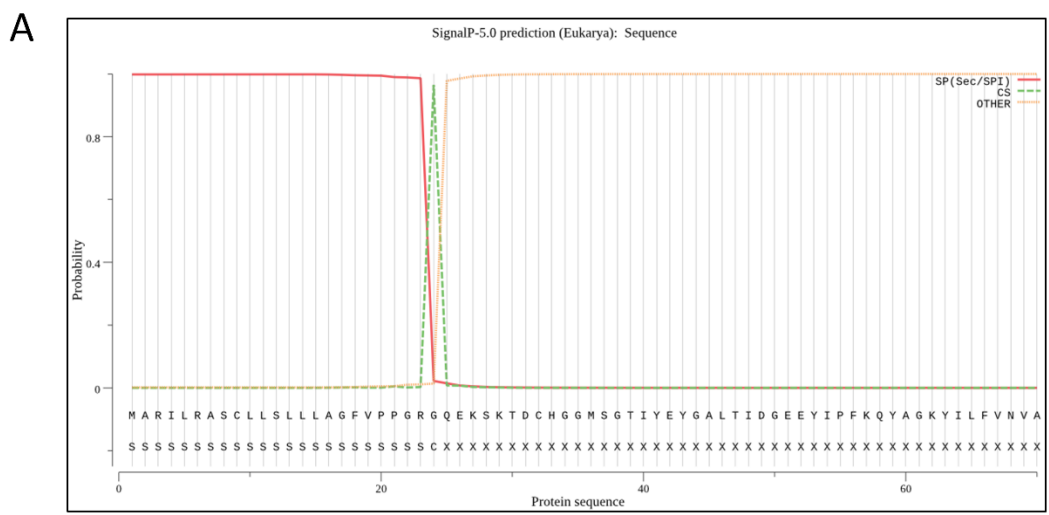

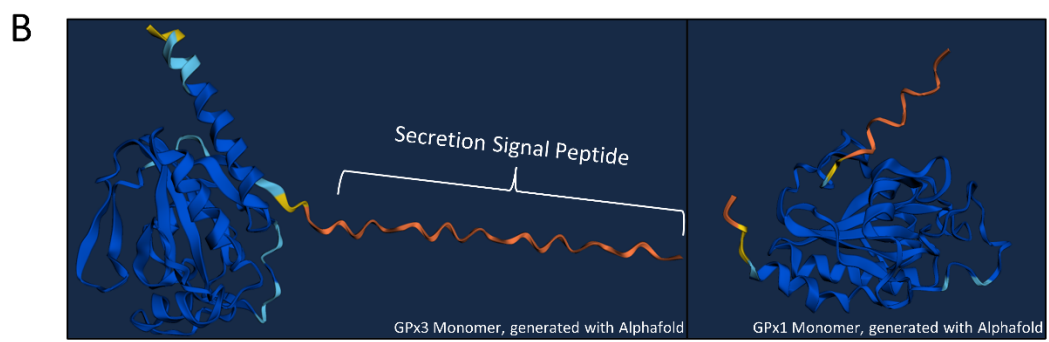

**Supplementary Figure S4:** GPX3 contains an N-terminal secretion signal peptide. **(A)** Probability of the presence of a secretion signal peptide within the N-terminal amino acid sequence of murine GPX3, determined by SignalP 5.0 (https://services.healthtech.dtu.dk/services/SignalP-5.0/) as described in Armenteros et al. (Nature biotech 2019). **(B)** Protein monomer structure prediction of full-length murine GPX3 (left panel) and GPX1 (right panel), using AlphaFold 3 (https://alphafold.ebi.ac.uk/).

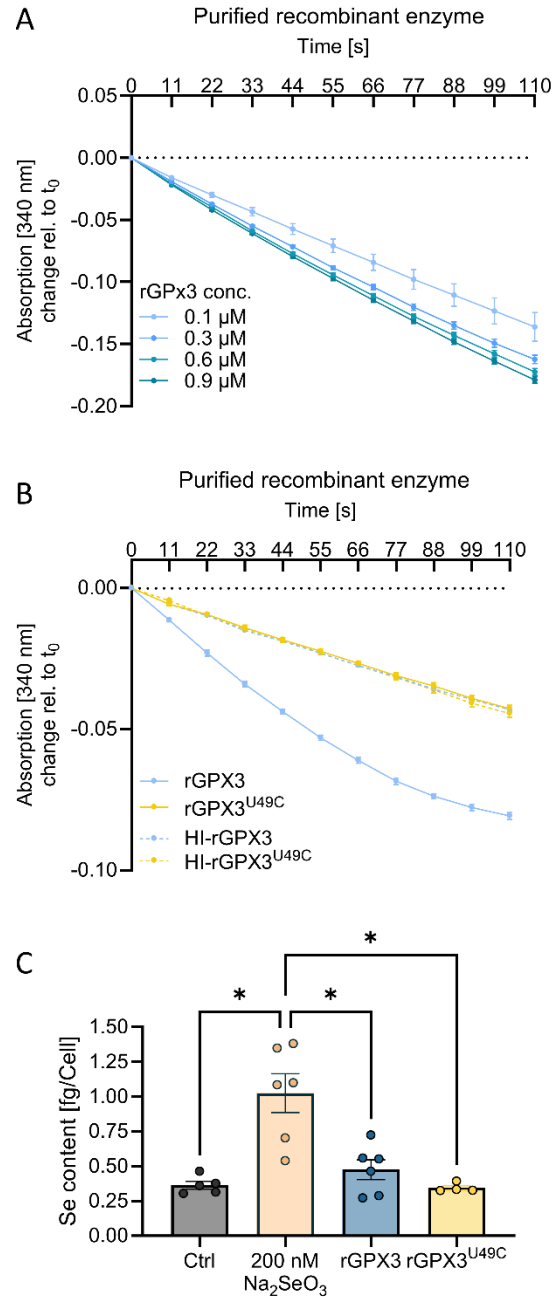

**Supplementary Figure S5:** Recombinant GPX3 shows dose-dependent activity, GPX3 does not serve as a selenium donor. **(A)** GPX activity assay of pure recombinant GPX3 in increasing concentrations, decrease in NADPH absorption signifies activity. **(B)** GPX activity assay of pure recombinant GPX3/rGPX3<sup>U49C</sup> and their heat-inactivated (HI) forms. **(C)** Total selenium content normalized to cell number after after sodium selenite (72 h 200 nM $\text{Na}_2\text{SeO}_3$), or rGPX3/rGPX3<sup>U49C</sup> treatment (D, 24 h, 500 ng/ml). All data are presented as mean ± SEM. Significances after 2-tailed Welch's t-test, \*: $p < 0.05$.

**Supplementary table S1: Amino acid sequences used for cloning the recombinant GPX3 enzymes.**
Position of selenocysteine and selenocysteine-coding sequence shown in bold and underlined.

| Protein | Synthetic DNA sequence | Protein sequence |
| --- | --- | --- |
| Mouse GPX3 without signal sequence | CAGGAGAAAAGTAAGACAGACTGCCACGGC<br>GGCATGTCAGGCACTATTTACGAGTATGGA<br>GCATTGACGATTGATGGCGAAGAATACATT<br>CCTTTCAAGCAATACGCTGGTAAGTACATTC<br>TTTTTGTAAACGTGGCCTCTTAT<u>**TAG**</u>GGTCT<br>GACGGATCAATACTTGGAATTGAATGCCTTG<br>CAAGAAGAATTGGGTCCATTTCGGACTTGTG<br>ATTTTAGGGTTCCTAGTAACCAGTTCGGTA<br>AACAGGAGCCCCGGCGAAAATAGCGAAATCT<br>TGCCCAGCTTGAAATACGTCCGTCCCGGAG<br>GCGGATTCGTGCCGAACCTCCAGTTATTCGA<br>AAAAGGGGATGTCAACGGAGAGAAGGAGC<br>AGAAATTCTATACTTTTCTGAAAAATAGTTG<br>CCCACCTACAGCGGAATTGTTAGGATCTCCC<br>GGTCGTCTGTTCTGGGAGCCGATGAAGATT<br>CACGACATTCGTTGGAACCTCGAGAAATTCC<br>TGGTCGGGCCCCGATGGCATTCCCGTGATGC<br>GCTGGTACCACCGTACTACGGTCAGTAACGT<br>GAAGATGGATATTTTGTCTATATGCGCCGT<br>CAAGCAGCTCTTCCGCTCGCGGCAAG | QEKS~~K~~TDCHGGMSGTIYEYGALTIDGEEYIPFKQ<br>YAGKYILFVNVASY<u>**U**</u>GLTDQYLELNALQEELGPF<br>GLVILGFPSNQFGKQEPGENSEILPSLKYVRPGG<br>GFVPNFQLFEKGDVNGEKEQKFYTLKNSCPPT<br>AELLGSPGR~~L~~FWEPMKIHDIRWNFEKFLVGPD<br>GIPVMRWYHRTTVSNVKMDILSYMRRQAALS<br>ARGK |
| Mouse GPX3 U49C without signal sequence | CAGGAGAAAAGTAAGACAGACTGCCACGGC<br>GGCATGTCAGGCACTATTTACGAGTATGGA<br>GCATTGACGATTGATGGCGAAGAATACATT<br>CCTTTCAAGCAATACGCTGGTAAGTACATTC<br>TTTTGTAAACGTGGCCTCTTAT<u>**TGT**</u>GGTCT<br>GACGGATCAATACTTGGAATTGAATGCCTTG<br>CAAGAAGAATTGGGTCCATTTCGGACTTGTG<br>ATTTTAGGGTTCCTAGTAACCAGTTCGGTA<br>AACAGGAGCCCCGGCGAAAATAGCGAAATCT<br>TGCCCAGCTTGAAATACGTCCGTCCCGGAG<br>GCGGATTCGTGCCGAACCTCCAGTTATTCGA<br>AAAAGGGGATGTCAACGGAGAGAAGGAGC<br>AGAAATTCTATACTTTTCTGAAAAATAGTTG<br>CCCACCTACAGCGGAATTGTTAGGATCTCCC<br>GGTCGTCTGTTCTGGGAGCCGATGAAGATT<br>CACGACATTCGTTGGAACCTCGAGAAATTCC<br>TGGTCGGGCCCCGATGGCATTCCCGTGATGC<br>GCTGGTACCACCGTACTACGGTCAGTAACGT<br>GAAGATGGATATTTTGTCTATATGCGCCGT<br>CAAGCAGCTCTTCCGCTCGCGGCAAG | QEKS~~K~~TDCHGGMSGTIYEYGALTIDGEEYIPFKQ<br>YAGKYILFVNVASY<u>**C**</u>GLTDQYLELNALQEELGPF<br>GLVILGFPSNQFGKQEPGENSEILPSLKYVRPGG<br>GFVPNFQLFEKGDVNGEKEQKFYTLKNSCPPT<br>AELLGSPGR~~L~~FWEPMKIHDIRWNFEKFLVGPD<br>GIPVMRWYHRTTVSNVKMDILSYMRRQAALS<br>ARGK |

**Supplementary table S2: Real-time quantitative PCR primer pairs**

| Target gene | Forward sequence | Reverse sequence |
| --- | --- | --- |
| *Beta-Actin* | GCCAACCGTGAAAAGATGAC | TACGACCAGAGGCATACAG |
| *Insr* | AATGGCAACATCACACACTACC | AATGGCAACATCACACACTACC |
| *GPX1* | ACAGTCCACCGTGTATGCCTTC | CTCTTCATTCTTGCCATTCTCCTG |
| *GPX3* | CCATTTGGCTTGGTCATTCTGGG | CACCTGGTCGAACATACTTGAGAC |
| *Ppary* | GCATGGTGCCTTCGCTGA | TGGCATCTCTGTGTCAACCATG |
| *C/ebpα* | CAAGAACAGCAACGAGTACCG | GTCACTGGTCAACTCCAGCAC |
| *Ap2* | AAGGTGAAGAGCATCATAACCCT | TCACGCCTTTCATAACACATTCC |
| *Plin1* | GACACACCATGCAAACCACAGCAT | TCGTCATGGCTCTCATCCTGA |
| *Glut4* | CATTCCCTGGTTCATTGTGG | GAAGACGTAAGGACCCATAGC |

**Supplementary table S3: Primary and secondary antibodies used for Western Blotting**

| Target | Species | Vendor | Cat # | RRID |
| --- | --- | --- | --- | --- |
| AKT | Mouse | Cell Signaling Technology | 2920 | AB_329827 |
| AKT<sub>S473</sub> | Rabbit | Cell Signaling Technology | 9271 | AB_329825 |
| GPX1 | Rabbit | Abcam | ab22604 | AB_2112120 |
| GPX3 | Rabbit | Proteintech | 13947-1-AP | AB_3085426 |
| InsR | Rabbit | Cell Signaling Technology | 3025 | AB_2280448 |
| InsR<sub>Y1150/1151</sub> | Mouse | Santa Cruz Biotechnology | sc-81500 | 1125642 |
| DNP<br>(Carbonylation) | Rabbit | Sigma-Aldrich | D9656 | AB_259258 |
| Anti-rabbit HRP | Goat | Cell Signaling Technology | 7074 | AB_2099233 |
| Anti-mouse HRP | Horse | Cell Signaling Technology | 7076 | AB_330924 |

**Supplementary table S4: Primer used for chromatin immunoprecipitation qPCR**

| Target gene | Forward sequence | Reverse sequence |
| --- | --- | --- |
| *Putative SP1 binding site* | GGTCTCTGAAGTTGCGGGTT | GGAGTCTGAGACAGAGGGACG |
| *Insulin gene* | CTTCAGCCCAGTTGACCAAT | AGGGAGGAGGAAAGCAGAAC |

**Supplementary table S5: Trace element-specific experimental conditions and recovery rates for metal quantification via ICP-MS/MS.**

| Element | Collision gas | Mass shift | Recovery rate (%) | Limit of Quantification (µg/L) |
| --- | --- | --- | --- | --- |
| Mn | He | 55 → 55 | 113.9 ± 4.2 | 0.2 |
| Fe | He | 56 → 56 | 114.0 ± 5.6 | 17.5 |
| Cu | He | 63 → 63 | 112.9 ± 4.1 | 0.8 |
| Zn | He | 66 → 66 | 119.9 ± 5.6 | 4.6 |
| Se | O<sub>2</sub> | 80 → 96 | 108.7 ± 6.0 | 0.2 |
| Ge | He, O<sub>2</sub> | 72 → 72 | - | - |
| Rh | He, O<sub>2</sub> | 103 → 103 | - | - |

**Supplementary table S6: Experimental conditions for metal quantification via ICP-MS/MS.**

|  |  |
| --- | --- |
| Plasma forward power | 1550 W |
| Plasma gas | 15.0 L/min |
| Auxiliary gas flow | 0.90 mL/min |
| Nebulizer gas flow | 0.99 mL/min |
| Nebulizer type | MicroMist |
| Collision gas | 4.5 L/min (He), 30 % (O<sub>2</sub>) |
| Calibration range | 0.2 – 1000 µg/L (Fe, Cu, Zn)<br>0.01 – 50 µg/L (Mn, Se) |
